# Transient protein accumulation at the center of the T cell antigen presenting cell interface drives efficient IL-2 secretion

**DOI:** 10.1101/296616

**Authors:** Danielle J. Clark, Laura E. McMillan, Sin Lih Tan, Gaia Bellomo, Clémentine Massoué, Harry Thompson, Lidiya Mykhaylechko, Dominic Alibhai, Xiongtao Ruan, Kentner L. Singleton, Minna Du, Alan J. Hedges, Pamela L. Schwartzberg, Paul Verkade, Robert F. Murphy, Christoph Wülfing

## Abstract

Supramolecular signaling assemblies are of interest for their unique signaling properties. A µm scale signaling assembly, the central supramolecular signaling cluster (cSMAC), forms at the center of the interface of T cells activated by antigen presenting cells. We have determined that it is composed of multiple complexes of a supramolecular volume of up to 0.5µm^3^ and associated with extensive membrane undulations. To determine cSMAC function, we have systematically manipulated the localization of three adaptor proteins, LAT, SLP-76, and Grb2. cSMAC localization varied between the adaptors and was diminished upon blockade of the costimulatory receptor CD28 and deficiency of the signal amplifying kinase Itk. Reconstitution of cSMAC localization restored IL-2 secretion which is a key T cell effector function as dependent on reconstitution dynamics. Our data suggest that the cSMAC enhances early signaling by facilitating signaling interactions and attenuates signaling thereafter through sequestration of a more limited set of signaling intermediates.

## Introduction

T cell activation is governed by spatiotemporal organization of signal transduction across scales. At the nanoscale receptors form clusters of dozens of molecules that can coalesce into microclusters and are commonly associated with active forms of signaling intermediates (Boyle et al., 2011; Hu et al., 2016; Lillemeier et al., 2006; Sherman et al., 2011; Varma et al., 2006; Yokosuka et al., 2005). This association suggests that such receptor clusters mediate efficient T cell signaling. Larger, µm scale assemblies were first described at the center and periphery of T cells activated by antigen presenting cells (APC) for the TCR, PKCθ and LFA-1, talin, respectively, as central and peripheral supramolecular activation clusters (cSMAC and pSMAC) (Grakoui et al., 1999; Monks et al., 1998; Monks et al., 1997). µm scale of assemblies, in particular in the form of supramolecular protein complexes, provides unique biophysical and signaling properties (Banani et al., 2016; Li et al., 2012; Shin and Brangwynne, 2017). Supramolecular protein complexes play critical roles in viral sensing (Cai et al., 2014), inflammation (Franklin et al., 2014), embryonic development (Brangwynne et al., 2009), protein folding in cancer (Rodina et al., 2016), nuclear ubiquitinoylation (Marzahn et al., 2016), and chromatin compaction (Larson et al., 2017). Such complexes are readily observed by fluorescence microscopy, held together by a network of multivalent protein interactions and often have distinct phase properties (Banani et al., 2016; Li et al., 2012; Shin and Brangwynne, 2017). The cSMAC has many properties of such supramolecular protein complexes: It contains various multivalent signaling intermediates (Balagopalan et al., 2015), prominently LAT (linker of activation of T cells), components of this complex including LAT and PKCθ exchange with the remainder of the cell to a moderate extent and slowly (Roybal et al., 2015), and components of this complex can be assembled into supramolecular structures *in vitro* (Su et al., 2016). Therefore, understanding biophysical properties of the cSMAC and how it regulates T cell activation is of substantial importance.

cSMAC function is controversial despite decades of work. Limitations in investigating the cSMAC are that its properties are largely unresolved and that its composition and/or assembly have not been systematically manipulated inside live T cells. Based on association of cSMAC formation with T cell activation conditions, the cSMAC has been proposed to enhance T cell signaling, terminate it, not be related to signaling or only upon weak stimulation or at late time points (Cemerski et al., 2008; Freiberg et al., 2002; Grakoui et al., 1999; Lee et al., 2002; Monks et al., 1997). cSMAC formation is often associated with efficient T cell activation conditions, fitting with a role in enhancing T cell signaling. Accumulation of signaling intermediates at the T cell:APC interface center is substantially reduced by blockade of the costimulatory receptor CD28 (Singleton et al., 2009; Wülfing et al., 2002), in regulatory T cells (Zanin-Zhorov et al., 2010), during thymic selection (Ebert et al., 2008), or in the absence of the signal amplifying kinase Itk (IL-2 inducible T cell kinase) (Singleton et al., 2011). To determine cSMAC properties, we have used stimulated emission depletion (STED) super-resolution microscopy and correlative light electron microscopy (CLEM). The cSMAC was composed of multiple complexes of supramolecular dimensions and associated with extensive membrane undulations. To investigate cSMAC function, we have systematically manipulated the localization of three adaptor proteins in live primary T cells: LAT (Balagopalan et al., 2015) is an integral component of the cSMAC. SLP-76 (SH2 domain-containing leucocyte protein of 76 kD) (Koretzky et al., 2006) is associated with it only during the first minute of T cell activation (Roybal et al., 2015). Grb2 (growth factor receptor-bound 2) (Jang et al., 2009) association with the cSMAC is less prevalent (Roybal et al., 2015). Interface recruitment of all three adaptors was diminished upon attenuation of T cell activation by costimulation blockade and Itk-deficiency as was IL-2 secretion, a critical T cell effector function. By fusing these adaptors with various protein domains with a strong interface localization preference we brought them back to the interface under the attenuated T cell activation conditions and restored cSMAC formation. Such restoration enhanced IL-2 secretion but only when executed to the extent and with dynamics seen under full stimulus conditions.

## Results

### µm scale LAT accumulation at the center of the T cell APC interface is associated with efficient T cell activation

To investigate the function of µm scale protein accumulation at the center of the T cell:APC interface, we first cataloged protein localization events that were consistently associated with efficient T cell activation. We attenuated T cell activation through costimulation blockade (Singleton et al., 2009; Wülfing et al., 2002) and Itk deficiency. The 5C.C7 T cell receptor (TCR) recognizes the moth cytochrome C (MCC) 89-103 peptide presented by I-E^k^. In the restimulation of *in vitro* primed 5C.C7 T cells with CH27 B cell lymphoma APCs and MCC peptide IL-2 amounts in the supernatant were reduced upon blockade of the CD28 ligands CD80 and CD86 (‘costimulation blockade’) and in T cells from Itk knock out 5C.C7 TCR transgenic mice, in particular at lower peptide concentrations (Fig. 1A). Even at an MCC peptide concentration of 10 µM the level of IL-2 mRNA in T cells is significantly (p < 0.001) reduced to less than 50% upon costimulation blockade and Itk-deficiency (Fig. 1B). 10 µM MCC was used for the remainder of the study.

**Figure 1.**
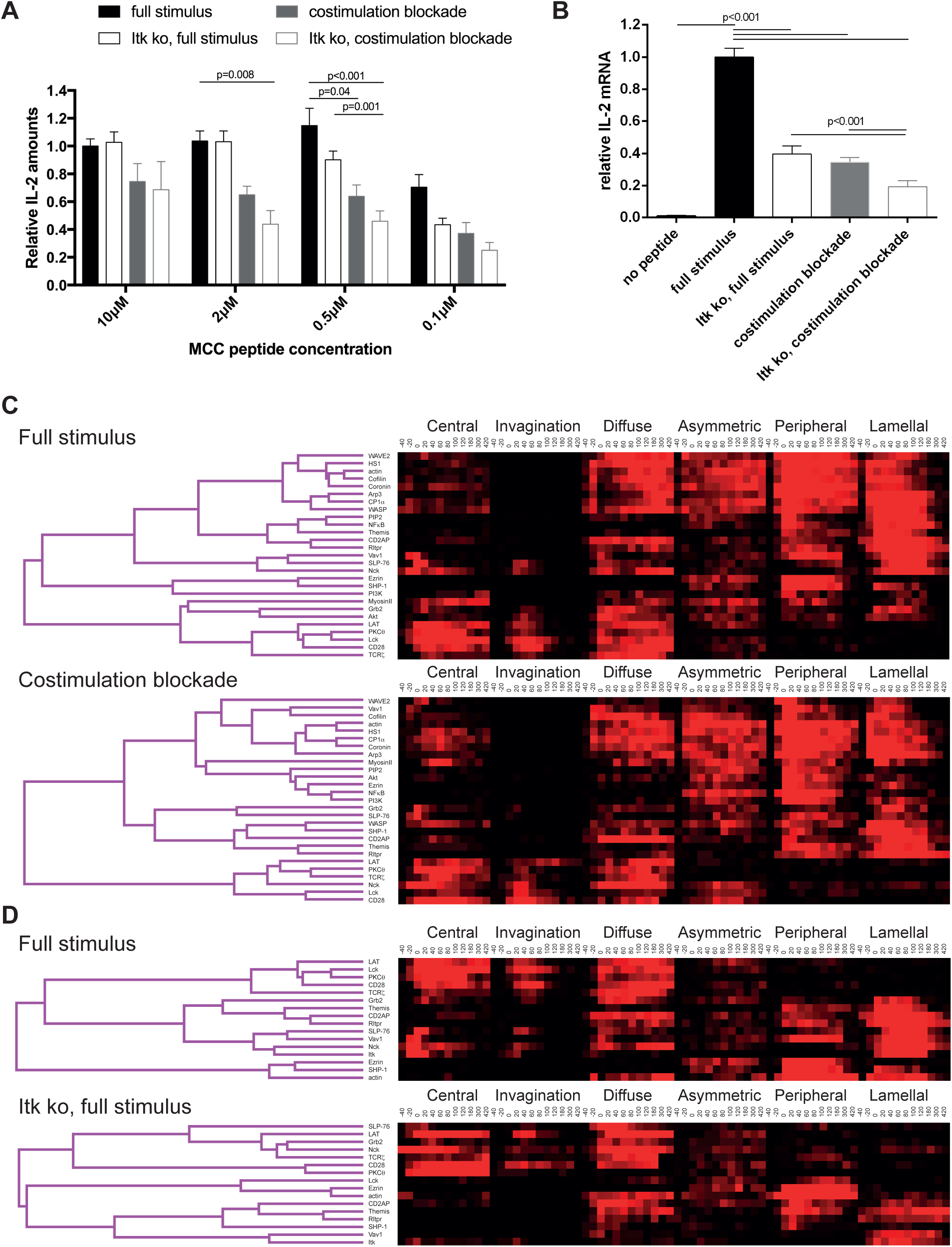
CD28 and Itk regulate IL-2 secretion and signaling organization. **A** *In vitro* primed 5C.C7 T cells, wild type or Itk-deficient (‘Itk ko’), were activated by CH27 APCs and the indicated concentration of MCC *pep*tide in the absence or presence of 10 µg/ml anti-CD80 plus anti-CD86 (‘full stimulus’ or ‘costimulation blockade’). IL-2 levels in the supernatant are given relative to stimulation of wild type 5C.C7 T cells under full stimulus conditions with 10 µM MCC with SEM. 4-8 experiments were averaged per condition. Statistical significance as determined separately for each MCC peptide concentration by 1-way ANOVA is indicated. **B** Relative levels of IL-2 mRNA are given upon 5C.C7 T cell activation similar to A with only 10 µM MCC. 3-18 experiments were averaged per condition. Statistical significance as determined by 1-way ANOVA is indicated. **C** 5C.C7 T cells expressing the indicated sensors were activated by CH27 B cell APCs (10 µM MCC) in the absence or presence of 10 µg/ml anti-CD80 plus anti-CD86 ((‘full stimulus’ or ‘costimulation blockade’) and percentage occurrence of patterns of interface enrichment (Fig. S1A) is given in shades of red from −40 to 420 s relative to tight cell coupling. Cluster trees are given in pink. **D** Similar to C pattern occurrence is given for wild type and Itk-deficient (‘Itk ko’) 5C.C7 T cells activated with a full stimulus. Some wild type 5C.C7 full stimulus data are the same as in C. Sensors used and source data for panels C and D are given in Table S1 and Figs. S1B-3.

We characterized T cell signaling organization as extensively described before (Ambler et al., 2017; Roybal et al., 2015; Singleton et al., 2009). Briefly, *in* vitro primed 5C.C7 T cells are retrovirally transduced to express fluorescent signaling intermediates or sensors, FACS sorted to low expression as close as possible to endogenous signaling intermediate concentrations and imaged in three dimensions over time during restimulation with APC and 10 µM MCC peptide (‘full stimulus’). In image analysis the frequency of occurrence of geometrically quantified µm scale subcellular distributions that represent underlying cell biological structures is determined (Fig. S1A) (Roybal et al., 2013). Of particular interest here are accumulation at the center of the T cell APC interface (‘central’), the cSMAC, and accumulation in a µm deep ‘invagination’ at the center of the interface that likely mediates termination of early central signaling (Singleton et al., 2006). Upon costimulation blockade and Itk deficiency the frequency of central accumulation of signaling sensors was consistently diminished (Fig. 1C, D, S1B-3, Table S1) indicative of reduced cSMAC formation. Efficient IL-2 secretion thus was associated with µm scale central signaling localization.

LAT as a key cSMAC component displayed µm scale central localization (Fig. 2A) in a biphasic pattern. At the time of tight cell coupling under full stimulus conditions 49 ± 6% of cell couples showed central LAT accumulation. After 2 min of cell coupling central LAT accumulation was only found in about 25% of cell couples (Fig. 2B) and remained stable at that level. In the absence of Itk, the initial peak of central LAT accumulation was significantly (p = 0.005) diminished to 26 ± 6% of cell couples with central LAT accumulation. Such reduction was more pronounced (11 ± 4%, p < 0.001) upon costimulation blockade (Fig. 2B, Table S2). Combining costimulation blockade and Itk deficiency yielded the least interface LAT accumulation (Fig. 2B). Impaired activation of cytoskeletal transport processes is a likely contributor to diminished central LAT accumulation upon costimulation blockade and Itk deficiency, as enhancement of actin dynamics with active Rac and Cofilin (Roybal et al., 2016) significantly (p < 0.001, Fig. 2C, Table S2)(Roybal et al., 2016) increased central and overall LAT accumulation.

**Figure 2.**
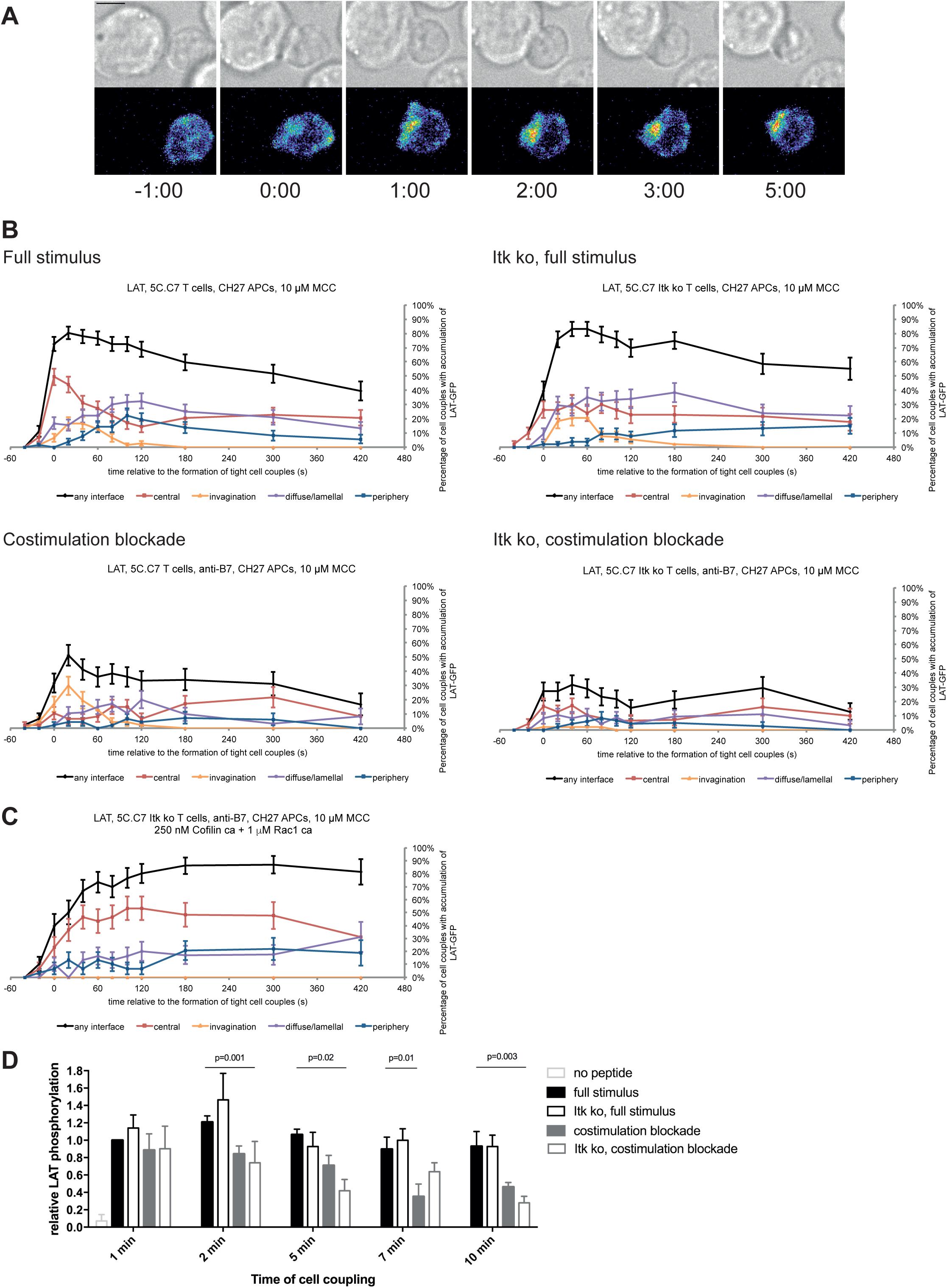
LAT localization and activation is regulated by costimulation and Itk. **A** An interaction of a LAT-GFP-transduced 5C.C7 T cell with a CH27 APC (10 μM MCC) is shown at the indicated time points (in minutes) relative to the time of formation of a tight cell couple. Differential interference contrast (DIC) images are shown in the top row, with top-down, maximum projections of 3-dimensional LAT-GFP fluorescence data in the bottom row. LAT-GFP fluorescence intensities are displayed in a rainbow-like false-color scale (increasing from blue to red). The scale bar corresponds to 5 µm. A corresponding video is available as video S1. **B** The graphs display the percentage of cell couples with LAT accumulation in the indicated patterns (Fig. S1A, ‘periphery’ is the sum of asymmetric and peripheral) relative to tight cell couple formation for wild type or Itk-deficient 5C.C7 T cells activated with CH27 APCs (10 µM MCC) in the absence or presence of 10 µg/ml anti-CD80 plus anti-CD86 (‘costimulation blockade’) as indicated. 47-77 cell couples were analyzed per condition, 226 total. A statistical analysis is given in Table S2. **C** Itk-deficient 5C.C7 T cells were activated with CH27 APCs (10 µM MCC) in the presence of 10 µg/ml anti-CD80 plus anti-CD86 and 250nM constitutively active Cofilin plus 1 µM constitutively active Rac1 as protein transduction reagents. LAT interface accumulation is given as in B. 30 cell couples were analyzed. Statistical significance is given in Table S2. **D** 5C.C7 T cells were activated as in B for the indicated times. Band intensities of α-LAT pY191 blots (Fig. S4) as normalized to the 1 min time point under full stimulus conditions are given. 5-7 experiments were averaged per condition. Statistical significance as determined separately for each time point by 1-way ANOVA is indicated.

Attenuation of T cell activation was associated with diminished LAT phosphorylation at Y191 (Fig. 2D, S4) upon costimulation blockade, in particular in combination with Itk deficiency. At 2, 5, and 10 min after tight cell coupling LAT phosphorylation was significantly (p ≤ 0.02) reduced in Itk-deficient 5C.C7 T cells upon costimulation compared to wild type 5C.C7 T cells under full stimulus conditions by 39%, 61%, and 70%, respectively.

### The cSMAC consists of multiple complexes of supramolecular dimensions and is associated with extensive membrane undulations

To determine cSMAC properties, we stained 5C.C7 T cell:CH27 B cell APC couples for LAT and LAT phosphorylated at Y191 (‘pLAT’) and imaged them using STED super-resolution microscopy (Fig. 3A). Multiple LAT/pLAT complexes formed in the cSMAC and/or beyond, on average 4 per cell. LAT complexes were significantly larger (p = 0.04) in the cSMAC region with a supramolecular volume of 0.23 ± 0.03µm^3^ than in T cells without a cSMAC (0.12 ± 0.01µm^3^)(Fig. 3B). pLAT complexes were similar with volumes of 0.25 ± 0.03µm^3^ (cSMAC) versus 0.13 ± 0.01µm^3^ (non-cSMAC, p = 0.007)(Fig. 3B).

**Figure 3.**
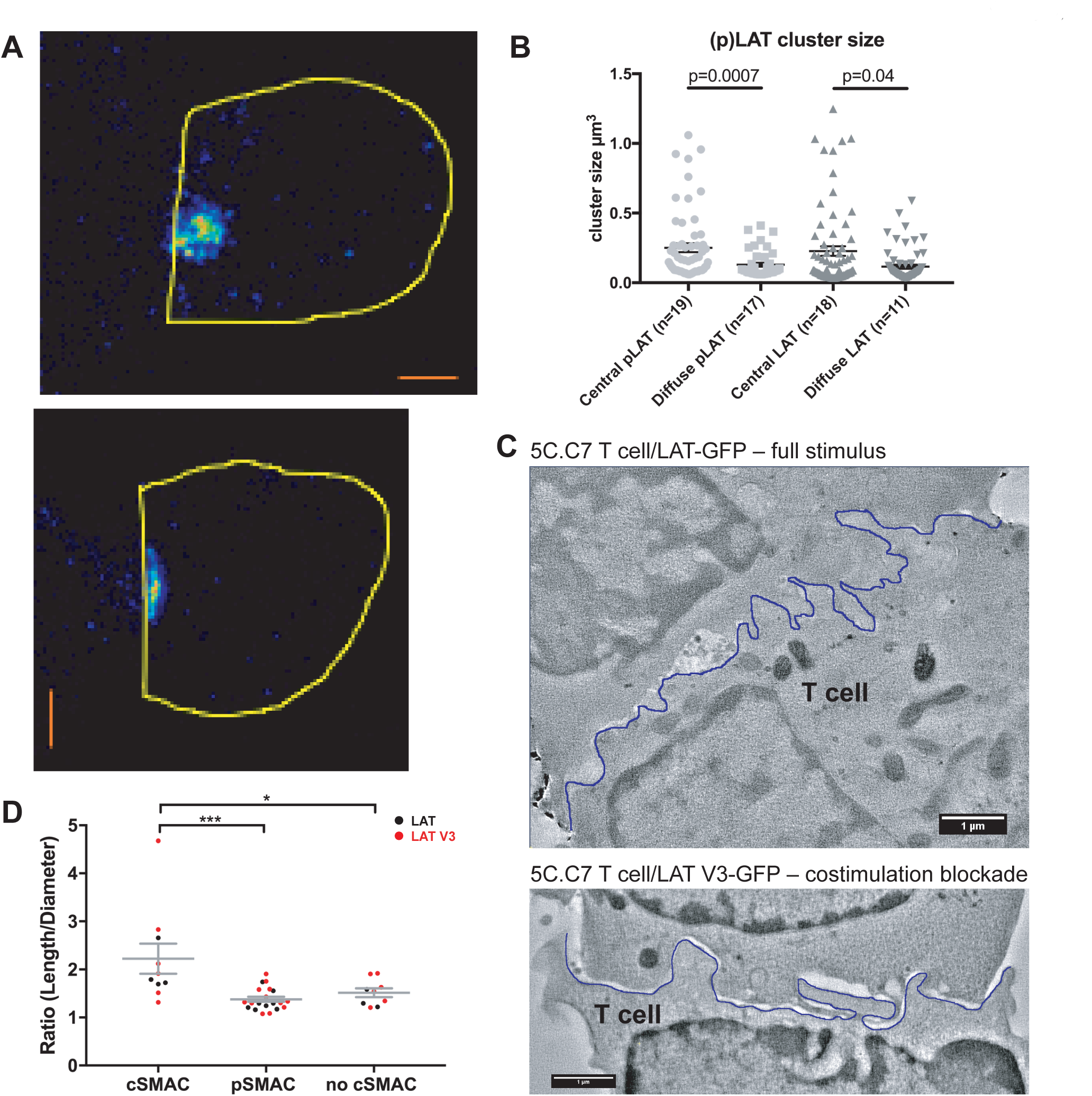
The cSMAC consists of multiple smaller complexes and is associated with extensive membrane undulations. **A** Two representative STED midplane images are given of 5C.C7 T cells activated by CH27 APCs (10 µm MCC) for 4.5 min and stained with α-LAT pY191. Staining fluorescence intensity is given in rainbow-like false-color scale (increasing from blue to red). The T cell outline is given in yellow. The scale bars correspond to 1 µm. **B** For experiments as in A LAT and LAT pY191 cluster size is given separately for cell couples with central or diffuse LAT accumulation as indicated (number of cell couples analysed in two independent experiments in parentheses). Statistical significance as determined separately for ‘LAT’ and ‘pLAT’ by Student’s test is indicated. **C** Midplane sections are given for two EM tomograms from a CLEM experiment. 5C.C7 T cells expressing LAT-GFP (top) or LAT V3-GFP (bottom) were activated by CH27 APCs under full stimulus (top) or costimulation blocked (bottom) conditions, respectively. Upon formation of a cell couple with central LAT-GFP or LAT V3-GFP accumulation cell were fixed and processed for EM. The T cell plasma membrane at the cellular interface is traced in blue. Videos of the entire EM tomogram reconstructions are given as videos S2, S3. **D** In cell couples processed as in C membrane undulations were determined as the ratio of the length of the plasma membrane (‘length’) to a straight-line interface diameter of the same region (‘diameter) in single images of EM sections. In cell couples with central LAT-GFP (black symbols) or LAT V3-GFP (red symbols) accumulation the interface center (‘cSMAC’) and periphery (‘pSMAC’) were measured separately. Peripheral regions were measured twice per cell, once to the left and once to the right of the central region. In control cell couples without central LAT-GFP (black symbols) or LAT V3-GFP (red symbols) accumulation (‘no cSMAC’) the entire interface was analyzed. Cell couples are derived from two independent experiments per condition. Statistical significance as determined by 1-way ANOVA is indicated (* p < 0.05, *** p < 0.001).

To understand how LAT as a transmembrane protein can drive the formation of multiple supramolecular complexes, we related cSMAC formation to plasma membrane topology with CLEM (Fig. 3C). During live cell imaging of 5C.C7 T cells activated by CH27 B cell APCs we rapidly fixed samples upon detection of central LAT clustering and processed them for electron microscopy (EM). In EM sections we measured the extent of membrane undulations in cSMAC and non-cSMAC interface regions as the ratio of the plasma membrane length to the straight-line diameter of the region. This ratio was significantly (p < 0.001) larger in the cSMAC (2.2 ± 0.3) than in the pSMAC of the same T cell (1.4 ± 0.1). In control cell couples without cSMAC formation the ratio as measured across the entire interface was as small (1.5 ± 0.1) as that in pSMAC regions of cell couples with central LAT clustering (Fig. 3D).

The cSMAC thus is a cellular region where membrane undulations and multiple supramolecular complexes drive enhanced proximity of signaling intermediates. To enable the throughput required for functional cSMAC investigation in the remainder of these studies, we used spinning disk confocal microscopy to determine central protein accumulation as a measure of cSMAC formation.

### Fusion of LAT with protein domains with pronounced interface localization preference controls LAT localization

To determine cSMAC function, we wanted to restore its formation upon costimulation blockade and in Itk-deficient T cells. To do so, we needed to hypothesize how cSMAC components interact within the complex. Conceptually, a supramolecular complex could function rigidly through formation of stoichiometrically defined protein interactions or flexibly by enhancing signaling proximity through complex formation such that the same proteins can be assembled using varying stoichiometries and protein interaction motifs. Because of the large number of cSMAC components some of which are membrane-bound (Fig. 1) we regard the flexible model as most likely. Impaired cSMAC formation upon costimulation blockade and Itk-deficiency should at least in part be driven by a reduction in the number of protein interaction motifs, i.e. the valence, of key components such as LAT as caused by diminished tyrosine phosphorylation (Fig. 2D). We should therefore be able to restore cSMAC formation by enhancing valence through the addition of new protein interaction domains. While this involves slightly divergent stoichiometries and protein interaction motifs the live cell signaling functionality gained through the formation of the central region of increased protein density should be restored.

We increased LAT valence by adding three protein domains: PKCθ V3, Vav1 SH3SH2SH3, or PLCδPH. The PKCθ V3 domain is required for central interface accumulation of PKCθ (Kong et al., 2011) even though it couldn’t drive central localization by itself (Fig. S5A). The Vav1 SH3SH2SH3 domains drove strong central accumulation only within the first minute of cell coupling (Fig. S5A), as consistent with the localization of full length Vav1 (Fig. S1B). The PLCδ PH domain mediated interface accumulation focused on the first two minutes of cell coupling without a central preference (Fig. S5A).

Fusion of LAT to the PKCθ V3 domain (LAT V3) yielded efficient central accumulation that was well sustained over the entire imaging time frame under all conditions at levels around 50% of cell couples with central accumulation (Fig. 4). Starting 40s after tight cell coupling such sustained central accumulation was significantly (p < 0.05, Table S3) more frequent than central accumulation of non-targeted LAT under full stimulus at almost every time point. Fusion of LAT to the PKCθ V3 domain thus stabilized central LAT accumulation well beyond the levels seen for LAT alone under any physiological condition. Nevertheless, in cSMAC formed upon LAT V3 expression membrane undulations were similarly enhanced as in T cells expressing LAT (Fig. 3D) supporting comparable cSMAC properties.

**Figure 4.**
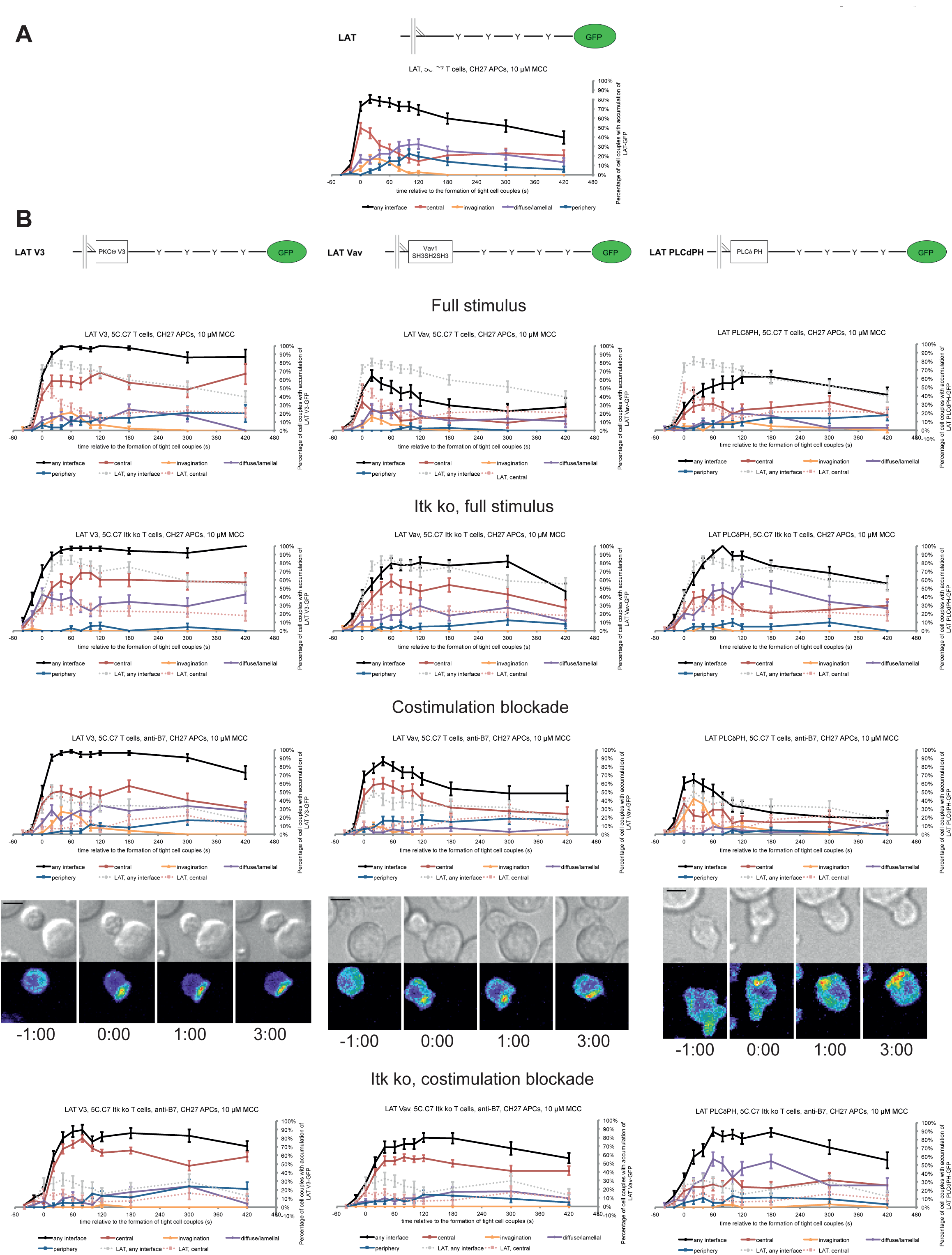
LAT localization can be controlled by fusion with protein domains with a strong interface localization preference. **A** A schematic representation of LAT-GFP is given (top) with LAT accumulation data under full stimulus conditions (bottom, from Fig. 2B) as a reference for the rest of the figure. **B** On top schematic representations are given for the three fusion proteins of LAT with protein localization domains as indicated. Corresponding imaging data are given in the respective columns below: wild type or Itk-deficient 5C.C7 T cells transduced to express the spatially targeted LAT construct indicated on the top of the column were activated with CH27 APCs (10 µM MCC) in the absence or presence of 10 µg/ml anti-CD80 plus anti-CD86 (‘costimulation blockade’) with different T cell activation conditions given in separate rows as indicated. Each individual graph gives the percentage of cell couples that displayed accumulation of the spatially targeted LAT construct with the indicated patterns as in Fig. 2B relative to tight cell couple formation in solid colors. Broken grey and red lines indicate accumulation of non-targeted LAT-GFP in any or the central interface pattern, respectively, under the same T cell activation conditions (from Fig. 2B) for reference. For costimulation blocked conditions representative imaging data are given similar to Fig. 2A. Corresponding videos are available as videos S4-6. 37-53 cell couples were analyzed per condition, 551 total. Statistical analysis is given in Table S3.

Fusion of LAT to the Vav1 SH3SH2SH3 domains (‘LAT Vav’) yielded different effects depending on the T cell activation conditions. Upon a full T cell stimulus LAT Vav resulted in diminished central and overall accumulation compared to LAT alone that was significant (p < 0.05) across many time points (Fig. 4, Table S3), suggesting that LAT Vav does not enhance properly assembled signaling assemblies. Upon the three attenuated stimuli LAT Vav consistently enhanced central accumulation, most dramatically (p ≤ 0.001, at most time points) for costimulation blockade in wild type and Itk deficient cells (Fig. 4, Table S3). LAT Vav accumulation upon attenuated T cell stimulation in any pattern was largely indistinguishable from non-targeted LAT accumulation under full stimulus conditions (p > 0.05, Table S3) and central accumulation was moderately enhanced only between 1 and 3 minutes after tight cell coupling. Fusion of LAT to the Vav1 SH3SH2SH3 domain thus allowed for fairly close reconstitution of full stimulus-type LAT localization upon attenuated T cell stimulation.

Fusion of LAT to the PLCδ PH domain (‘LAT PLCδPH’) resembled LAT Vav but was less powerful (Fig. 4, Table S3). Upon the three attenuated T cell stimuli LAT PLCδPH moderately enhanced central and overall LAT accumulation but didn’t consistently reach the same extent as LAT alone under full stimulus conditions. For example, in Itk-deficient 5C.C7 T cells upon costimulation blockade interface accumulation in any pattern was consistently enhanced (p < 0.005 for time point 20 and later) from <32% to >55% upon expression of LAT PLCδPH. However, accumulation at the interface center only moderately increased from a range of 6-17% to 19-35%.

To ensure that overall interface accumulation of the targeted LAT constructs was comparable, we measured (Ambler et al., 2017; Roybal et al., 2016) their interface recruitment upon costimulation blocked conditions. All constructs showed substantial interface recruitment with moderately less LAT Vav recruitment at the last four time points (Fig. S5B, C). To ensure functionality of the targeted LAT constructs, we showed that they were tyrosine phosphorylated upon T cell activation (Fig. S5D).

Fusion of LAT with additional protein interaction domains thus allowed us to control central clustering: Fusion with the PKCθ V3 domain yielded consistently enhanced central localization, fusion with the Vav1 SH3SH2SH3 domains largely restored full stimulus-type LAT localization upon costimulation blockade and Itk deficiency and fusion with the PLCδ PH domain resulted in partial restoration.

### Restoration of LAT centrality yields enhanced IL-2 mRNA production

To determine T cell function upon manipulation of LAT valence and localization, we measured IL-2 mRNA induction upon 5C.C7 T cell activation with CH27 APCs and 10 µM MCC peptide, directly mirroring the imaging conditions. Expression of the targeting domains in isolation had only minor effects on IL-2 mRNA amounts (Fig. S6). Forcing exaggerated central LAT clustering by fusion with the PKCθ V3 domain did not affect IL-2 mRNA amounts (Fig. 5A, B). In contrast, restoring LAT centrality under costimulation blocked and Itk deficient conditions to slightly higher (LAT Vav) or slightly lower (LAT PLCδPH) levels than seen for non-targeted LAT under full stimulus conditions yielded a consistent and largely significant (p < 0.05) increase in IL-2 mRNA (Fig. 5A, B) to levels close to the amounts of IL-2 mRNA in LAT-transduced 5C.C7 T cells under full stimulus conditions. For example, in LAT-expressing 5C.C7 T cells IL-2 mRNA amounts dropped to 18 ± 6% and 18 ± 3% of full stimulus mRNA upon costimulation blockade in wild type and Itk-deficient 5C.C7 T cells, respectively. Expression of LAT Vav restored IL-2 mRNA to 41 ± 9% and 58 ± 10% (p < 0.01), respectively, and expression of LAT PLCδPH to 50 ± 9% (p < 0.05) and 88 ± 23%. µm scale central LAT interface accumulation thus supported efficient IL-2 secretion depending on accumulation extent and dynamics.

**Figure 5.**
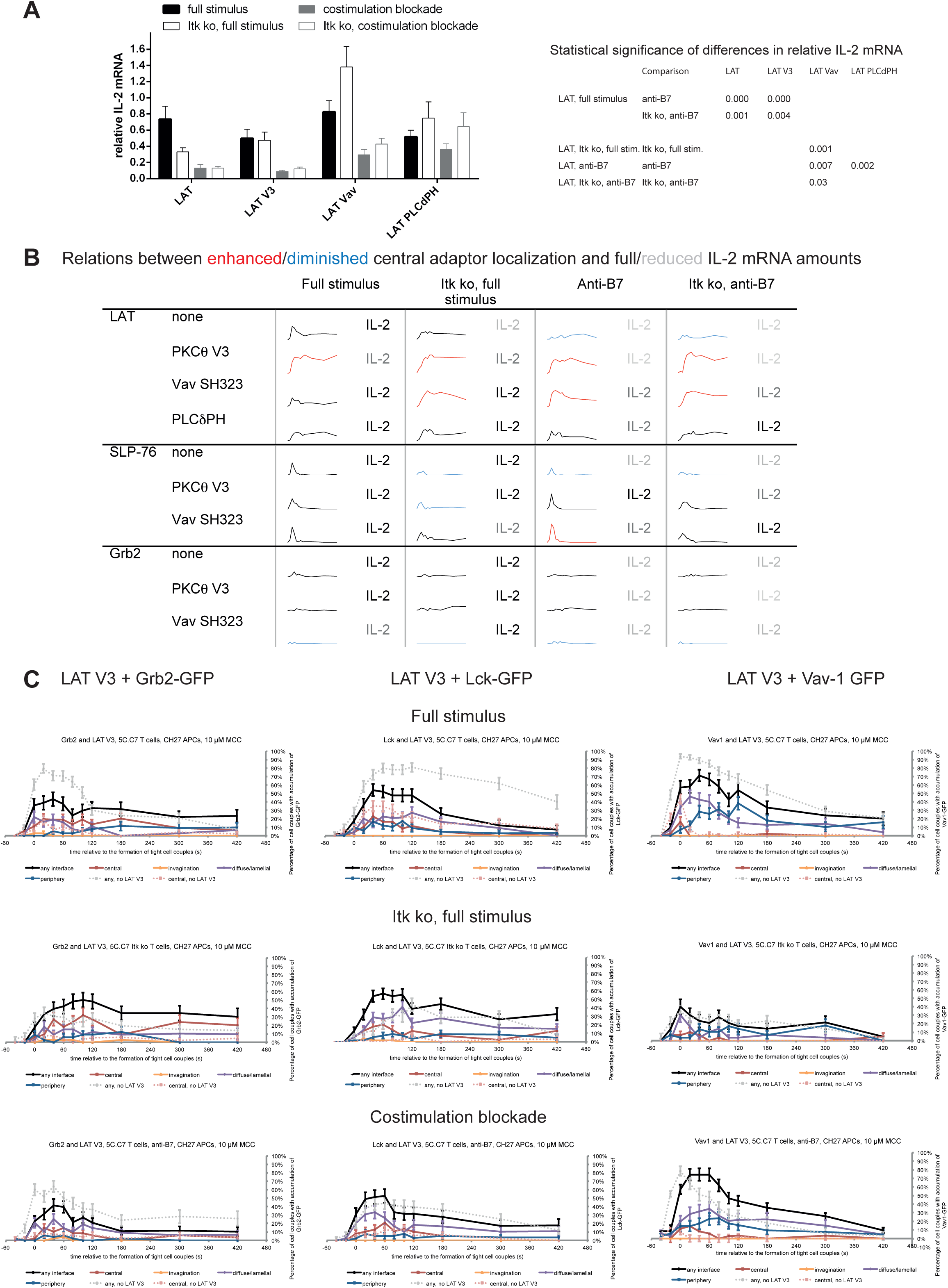
Restoration of LAT centrality enhances IL-2 generation but only modestly affects centrality of other signaling intermediates. **A** Wild type or Itk-deficient (‘Itk ko’) 5C.C7 T cells expressing LAT-GFP or a spatially targeted variant thereof as indicated were activated by CH27 APCs (10 µM MCC) in the absence or presence of 10 µg/ml anti-CD80 plus anti-CD86 (‘costimulation blockade’). IL-2 mRNA amounts are given relative to IL-2 mRNA in non-transduced 5C.C7 T cells under full stimulus conditions. 3-9 experiments were averaged per condition. Statistical significance as determined by 2-way ANOVA is given in the table on the right. **B** Sensor accumulation at the interface center and IL-2 mRNA amounts are summarized. Traces are the percentage cell couples with central accumulation from Figs. 4, 6, 7. Red indicates enhanced central accumulation relative to non-targeted signaling intermediate under full stimulus conditions at at least half of the time points with substantial central accumulation, blue similarly indicates diminished central accumulation. Four shades of grey indicated level of IL-2 mRNA relative to non-targeted signaling intermediate under full stimulus conditions >75%, 50-75%, 25-50% and <25%. **C** Wild type and Itk-deficient 5C.C7 T cells were transduced to express LAT V3 together with the indicated GFP-tagged signaling intermediate and activated with CH27 APCs (10 µM MCC) in the absence or presence of 10 µg/ml anti-CD80 plus anti-CD86 (‘costimulation blockade’) with different T cell activation conditions given in separate rows as indicated. Each individual graph gives the percentage of cell couples that displayed accumulation of the GFP-tagged signaling intermediate with the indicated patterns as in Fig. 2B relative to tight cell couple formation in solid colors. Broken grey and red lines indicate accumulation of the signaling intermediate in the absence of LAT V3 in any or the central interface pattern, respectively, under the same T cell activation conditions (from Fig. 7, Figs. S1B-3). 30-56 cell couples were analyzed per condition, 373 total. Statistical analysis is given in Table S4.

### Forcing central LAT localization only modestly enhances the central localization of related signaling intermediates

Next, we investigated to which extent the forced relocalization of one signaling intermediate can drive analogous relocalization of others. We determined the subcellular distributions of Grb2, Lck and Vav1 in 5C.C7 T cells in the presence of LAT V3 using IRES-containing retroviral vectors for the parallel expression of GFP-tagged versions of the signaling intermediates alongside LAT V3. Under full stimulus conditions expression of LAT V3 moderately diminished interface recruitment of Grb2, Lck and Vav1 (Fig. 5C, Table S4) suggesting that excessive central LAT localization upsets a finely balanced signaling system. Upon costimulation blockade the localization of all three signaling intermediates was largely unaffected by LAT V3. However, Grb2 and Lck centrality was moderately enhanced in Itk-deficient 5C.C7 T cells. For example, while the percentage of Itk-deficient 5C.C7 T cells with central Grb2-GFP expression did not exceed 7% at 40s after cell coupling and thereafter, upon co-expression of LAT V3 this percentage averaged 15% over the same time frame. The centrality of Vav1 as a signaling intermediate with only moderate early central accumulation (Fig. S1B) was not altered. The forced central localization of LAT thus could only draw in Grb2 and Lck as signaling intermediates with some intrinsic central localization preference to a modest extent and upon only one attenuated T cell stimulus.

### Restoration of SLP-76 centrality modestly enhances IL-2 mRNA production

We investigated SLP-76 as an adaptor with more transient central accumulation. At the time of tight cell coupling under full stimulus conditions 45 ± 7% of the cell couples displayed SLP-76 accumulation at the interface center, similar to LAT. However, 80s later this percentage dropped to less than 10% (Fig. 6A, B). Also similar to LAT, the peak of central SLP-76 accumulation was significantly (p ≤ 0.01) diminished upon costimulation blockade, Itk-deficiency and the combination of both to <27%, <15% and <11% of cell couples with central SLP-76 accumulation, respectively (Fig. 6B, Table S5). To enhance SLP-76 valence and thus control its localization, we fused SLP-76 to the PKCθ V3 (‘SLP-76 V3’) or the Vav1 SH3SH2SH3 (‘SLP-76 Vav’) domain. Both constructs did not significantly affect SLP-76 centrality under full stimulus conditions, in wild type or Itk-deficient 5C.C7 T cells (Fig. 6C, Table S5). However, accumulation of SLP-76 at the interface center was moderately but significantly (p < 0.05 at at least two time points within the first minute of tight cell coupling, the peak of central SLP-76 accumulation) enhanced upon costimulation blockade in wild type and Itk-deficient 5C.C7 T cells reaching e.g. 67 ± 7% and 33 ± 7%, respectively, of cell couples with central SLP-76 accumulation upon expression of SLP-76 Vav (Fig. 6C, Table S5). Interestingly, the enhancement of SLP-76 centrality was limited to the first minute of tight cell coupling, the time where non-targeted SLP-76 accumulated at the interface center.

**Fig. 6.**
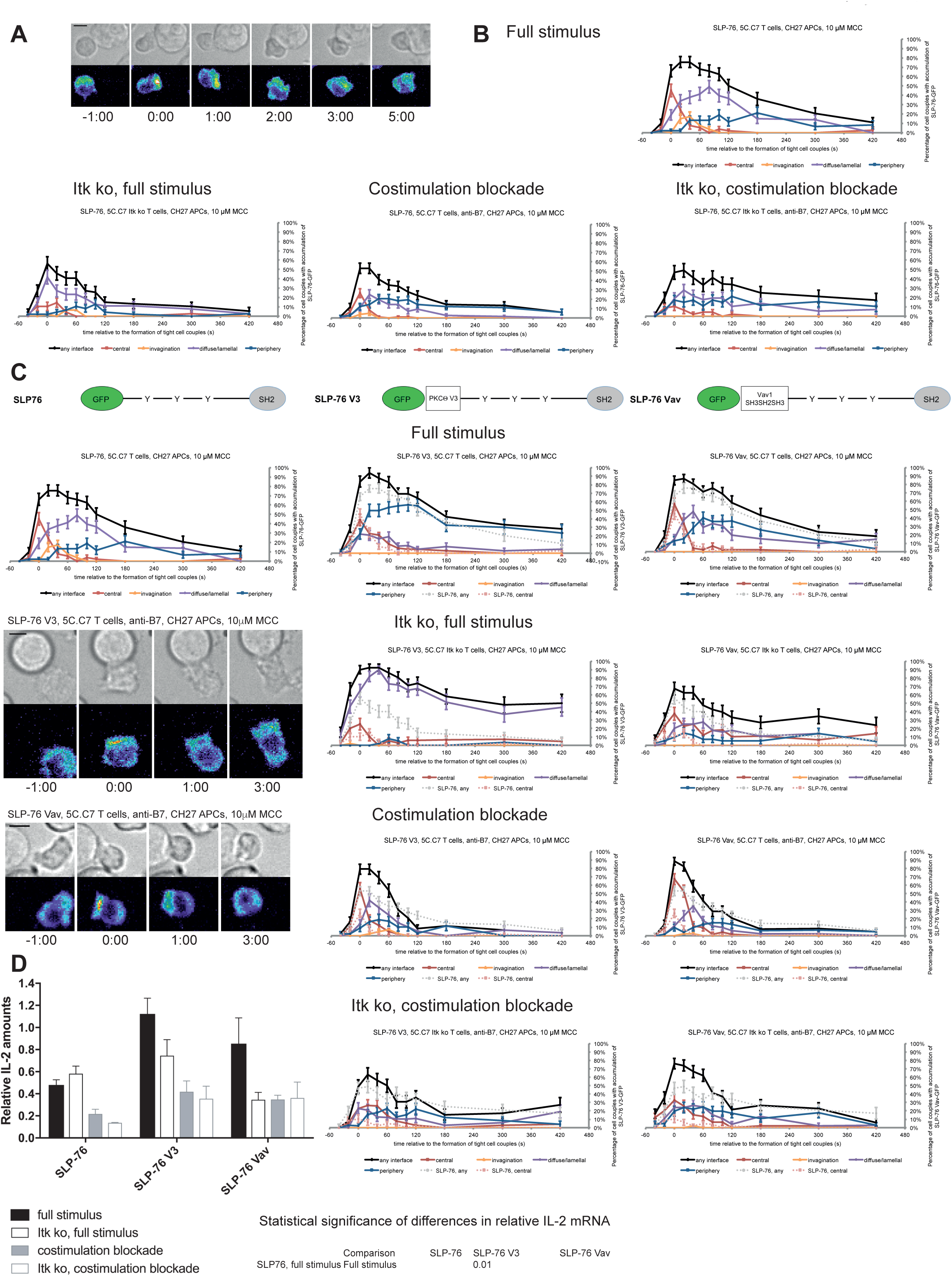
SLP-76 localization is regulated by costimulation and Itk and restoration of early SLP-76 centrality enhances IL-2 secretion. **A** An interaction of a SLP-76-GFP-transduced 5C.C7 T cell with a CH27 APC (10 μM MCC) is shown at the indicated time points (in minutes) relative to the time of formation of a tight cell couple as in Fig. 2A. A corresponding video is available as video S7. **B** The graphs display the percentage of cell couples with SLP-76 accumulation in the indicated patterns as in Fig. 2B relative to tight cell couple formation for wild type or Itk-deficient 5C.C7 T cells activated with CH27 APCs (10 µM MCC) in the absence or presence of 10 µg/ml anti-CD80 plus anti-CD86 (‘costimulation blockade’) as indicated. 47-83 cell couples were analyzed per condition, 231 total. A statistical analysis is given in Table S5. **C** On top schematic representations are given for SLP-76-GFP and the two fusion proteins of SLP-76 with protein domains as indicated. Corresponding imaging data are given in the respective columns below: wild type or Itk-deficient 5C.C7 T cells transduced to express the spatially targeted SLP-76 construct indicated on the top of the column were activated with CH27 B cell APCs (10 µM MCC) in the absence or presence of 10 µg/ml anti-CD80 plus anti-CD86 (‘costimulation blockade’) with different T cell activation conditions given in separate rows as indicated. Each individual graph gives the percentage of cell couples that displayed accumulation of the non-targeted (on the left for reference) or spatially targeted SLP-76 construct (middle and right) with the indicated patterns as in Fig. 2B relative to tight cell couple formation in solid colors. Broken grey and red lines indicate accumulation of non-targeted SLP-76-GFP in any or the central interface pattern, respectively, under the same T cell activation conditions (from B). For costimulation blocked conditions representative imaging data are given on the middle left similar to Fig. 2A. Corresponding videos are available as videos S8, 9. 39-52 cell couples were analyzed per condition, 355 total. Statistical analysis is given in Table S5. **D** Wild type or Itk-deficient (‘Itk ko’) 5C.C7 T cells expressing SLP-76-GFP or a spatially targeted variant thereof as indicated were activated by CH27 APCs (10 µM MCC) in the absence or presence of 10 µg/ml anti-CD80 plus anti-CD86 (‘costimulation blockade’). IL-2 mRNA amounts are given relative to IL-2 mRNA in non-transduced 5C.C7 T cells under full stimulus conditions. 2-8 experiments were averaged per condition. Statistical significance as determined by 2-way ANOVA is given on the bottom right.

Consistent with the enhancement of SLP-76 centrality we observed a modest increase in IL-2 mRNA amounts under costimulation blocked conditions upon expression of SLP-76 V3 and SLP-76 Vav. Upon expression of non-targeted SLP-76 costimulation blockade in wild type and Itk-deficient 5C.C7 T cells reduced IL-2 mRNA amounts to 45 ± 9% and 21 ± 1%, respectively, of full stimulus (Fig. 6D). Expression of SLP-76 V3 restored IL-2 mRNA amounts to 87 ± 20% and 74 ± 24%, respectively, expression of SLP-76 Vav to 73 ± 8% and 75 ± 30% without reaching statistical significance in the stringent 2-way ANOVA (Fig. 6D). Importantly, enhancement of centrality and IL-2 secretion remained closely linked across multiple T cell activation conditions and spatially targeted SLP-76 constructs (Fig. 5B) thus corroborating the importance of µm scale central protein accumulation within the first two minutes of cell coupling for IL-2 secretion.

### PKCθ V3 and Vav1 SH3SH2SH3 don’t affect Grb2 centrality and IL-2 mRNA production

As a negative control we enhanced the valence of a signaling intermediate with more tentative central localization preference: We fused Grb2 to PKCθ V3 or Vav1 SH3SH2SH3. Upon 5C.C7 T cell activation with a full stimulus Grb2 is efficiently recruited to the T cell:APC interface during the first two minutes of tight cell coupling, peaking at 80 ± 5% of cell couples with any interface accumulation. This overall interface accumulation was significantly (p ≤ 0.02 at at least three time points within the first two minutes of tight cell coupling) diminished upon costimulation blockade, Itk-deficiency and both, remaining below 64%, 35% and 43%, respectively of cell couples with any interface accumulation (Fig. 7A, B, Table S6). Distinguishing Grb2 from LAT and SLP-76, Grb2 accumulation at the interface center didn’t exceed 20% under any of the T cell activation conditions. Fusion of Grb2 with PKCθ V3 (‘Grb2 V3’) or Vav1 SH3SH2SH3 (‘Grb2 Vav’) did not enhance centrality under any of the T cell activation conditions (Fig. 7C, Table S6). Fusion with the PKCθ V3 domain did not substantially alter Grb2 localization at all (Fig. 7C). Fusion with the Vav1 SH3SH2SH3 domain enhanced overall Grb2 interface recruitment across many time points under all T cell activation conditions (Fig. 7C). However, most of this accumulation was in the peripheral pattern, a pattern common with full length Vav1 (Fig. S1B). As an important negative control, a protein with a minor intrinsic central preference can thus not be forced to the interface center. Expression of Grb2 V3 or Grb2 Vav did not alter IL-2 mRNA production under any of the T cell activation conditions (Fig. 7D) despite the substantially enhanced overall interface accumulation upon expression of Grb2 Vav. These data thus provide an important specificity control for the selective functional importance of protein accumulation in the cSMAC.

**Fig. 7.**
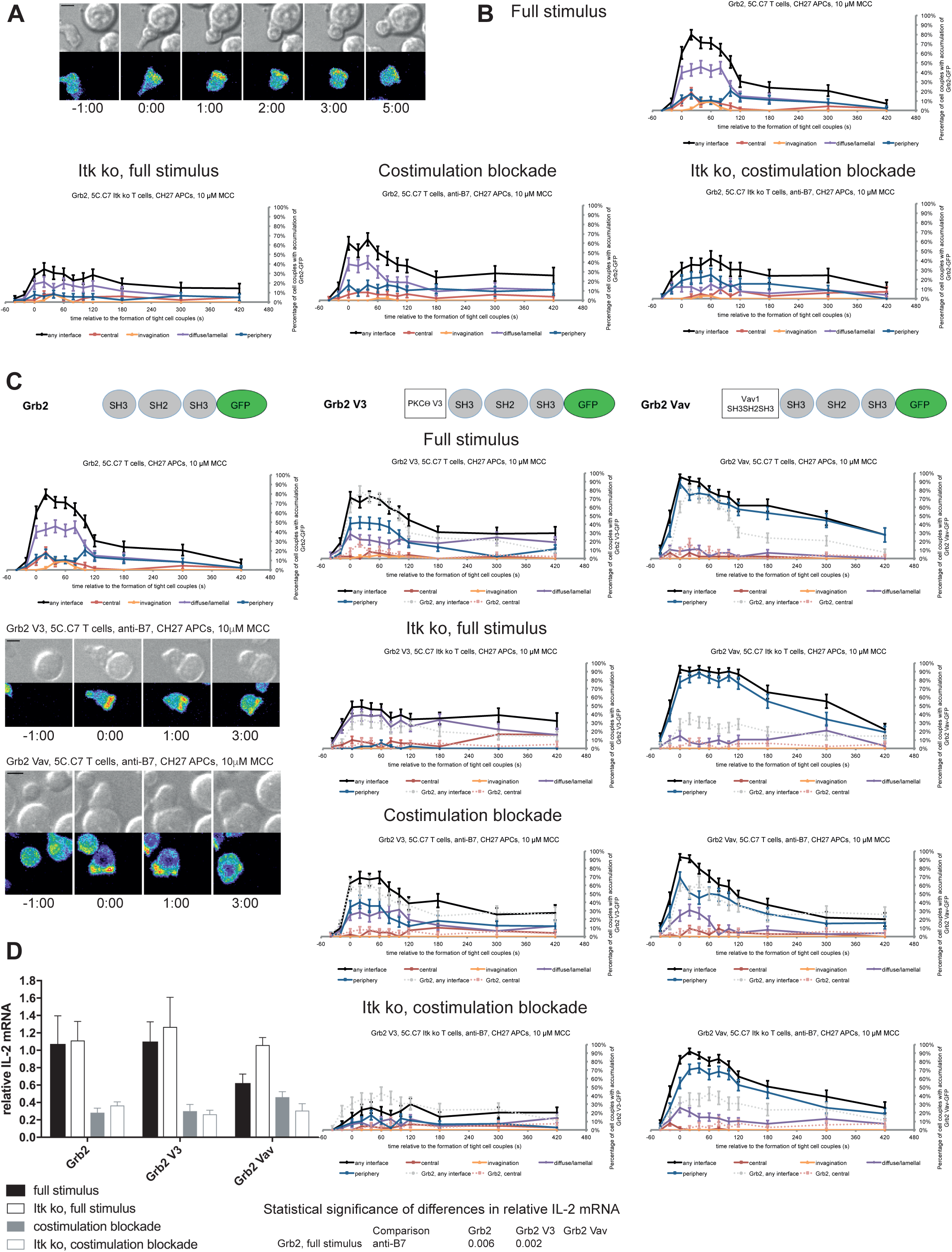
Grb2 localization is regulated by costimulation and Itk and doesn’t regulate IL-2 secretion. **A** An interaction of a Grb2-GFP-transduced 5C.C7 T cell with a CH27 APC (10 μM MCC) is shown at the indicated time points (in minutes) relative to the time of formation of a tight cell couple as in Fig. 2A. A corresponding video is available as video S10. **B** The graphs display the percentage of cell couples with Grb2 accumulation in the indicated patterns as in Fig. 2B relative to tight cell couple formation for wild type or Itk-deficient 5C.C7 T cells activated with CH27 APCs (10 µM MCC) in the absence or presence of 10 µg/ml anti-CD80 plus anti-CD86 (‘costimulation blockade’) as indicated. 42-59 cell couples were analyzed per condition, 204 total. A statistical analysis is given in Table S6. **C** On top schematic representations are given for Grb2-GFP and the two fusion proteins of Grb2 with protein domains as indicated. Corresponding imaging data are given in the respective columns below: wild type or Itk-deficient 5C.C7 T cells transduced to express the spatially targeted Grb2 construct indicated on the top of the column were activated with CH27 APCs (10 µM MCC) in the absence or presence of 10 µg/ml anti-CD80 plus anti-CD86 (‘costimulation blockade’) with different T cell activation conditions given in separate rows as indicated. Each individual graph gives the percentage of cell couples that displayed accumulation of the non-targeted (on the left for reference) or spatially targeted Grb2 construct (middle and right) with the indicated patterns as in Fig. 2B relative to tight cell couple formation in solid colors. Broken grey and red lines indicate accumulation of non-targeted Grb2-GFP in any or the central interface pattern, respectively, under the same T cell activation conditions (from B). For costimulation blocked conditions representative imaging data are given on the middle left similar to Fig. 2A. Corresponding videos are available as videos S11, 12. 41-62 cell couples were analyzed per condition, 387 total. Statistical analysis is given in Table S6. **D** Wild type or Itk-deficient (‘Itk ko’) 5C.C7 T cells expressing Grb2-GFP or a spatially targeted variant thereof as indicated were activated by CH27 APCs (10 µM MCC) in the absence or presence of 10 µg/ml anti-CD80 plus anti-CD86 (‘costimulation blockade’). IL-2 mRNA amounts are given relative to IL-2 mRNA in non-transduced 5C.C7 T cells under full stimulus conditions. 2-5 experiments were averaged per condition. Statistical significance as determined by 2-way ANOVA is given on the bottom right.

## Discussion

The cSMAC was characterized by enhanced membrane undulations and the formation of multiple protein complexes of supramolecular dimensions in the 0.1-0.5µm^3^ range. Supramolecular complexes can be driven by the polymerization of a single or few defined proteins (Cai et al., 2014; Franklin et al., 2014; Li et al., 2012; Marzahn et al., 2016) or, likely most common, they can consists of an agglomeration of large numbers of proteins (Rodina et al., 2016; Tarantino et al., 2014). While supramolecular complexes formed by a small number of components are often characterized by defined structures such as lipid droplets or fibers (Shin and Brangwynne, 2017), our understanding of supramolecular complexes built from a large number of components is limited. As close to half of the amount of three components of the central signaling complex, LAT, active Rac, and PKCθ, is immobile and the remainder exchanges only slowly with rest of the cell (Roybal et al., 2015) cSMAC signaling complexes likely have partial solid state properties. T cell signaling has been extensively characterized with super-molecular resolution in T cells activated with planar APC substitutes, supported lipid bilayers and antibody-coated cover slips. Similar to our findings large protein complexes, microclusters, were found. The microclusters are transported to the interface center to form a single µm scale structure (Varma et al., 2006; Yokosuka et al., 2005). However, in T cell:APC couples membrane undulations across the entire interface with F-actin structures perpendicular to the interface plane (Roybal et al., 2015) impair centripetal transport, a large central invagination may remove proteins from the interface center (Singleton et al., 2006) and the central membrane undulations characterized here increase the number of transmembrane proteins in contact with a given cytoplasmic volume. Therefore, the cSMAC in T cell:APC couples will likely have similar molecular constituents as the central complex in T cells activated on planar substrates but distinct biophysical properties. It is the signaling functionality arising from these unique properties that we have investigated here.

Using a large-scale live cell imaging approach, we were able to distinguish between inefficient cSMAC formation and inefficient recruitment of a signaling intermediate to an existing cSMAC: As central interface accumulation of numerous signaling intermediates was consistently diminished upon costimulation blockade and Itk-deficiency (Fig. 1C, D) cSMAC formation was most likely impaired. Central interface accumulation of spatially targeted LAT and SLP-76 upon the attenuated T cell stimuli thus has to represent cSMAC restoration as confirmed by CLEM (Fig. 3D).

Complexes of supramolecular dimensions are part of the cSMAC (Fig. 3B). In general, supramolecular complex formation becomes more likely with increasing concentrations and valences of the complex components (Banani et al., 2016; Li et al., 2012). Accordingly, replacement of proline-rich regions in Sos that mediate multivalent LAT/Grb2/Sos interactions leads to reduced LAT clustering and phosphorylation (Kortum et al., 2013). Similarly, when LAT phosphorylation (Fig. 2C), which is required for interactions of LAT with SH2 domain-containing signaling intermediates, was reduced upon costimulation blockade and Itk deficiency LAT clustering at the interface center was also diminished (Fig. 2 B). Compensation for such diminished LAT valence by fusing LAT with additional protein interaction domains restored µm scale central LAT accumulation. The dependence of LAT clustering on the number of functional protein interaction motifs, i.e. its valence, further supports the supramolecular nature of protein complexes within the cSMAC.

Interestingly, fusion domains could not drive spatial features of adaptor localization by themselves but could only enhance intrinsic adaptor localization preferences. We could enhance central LAT clustering across all time points, central SLP-76 clustering only during the first minute of cell coupling and central Grb2 clustering not at all. This mirrored the accumulation patterns of the non-targeted adaptors under full stimulus conditions (Figs. 4, 6, 7) and therefore strongly suggests that localization of the spatially targeted adaptors was driven by a combination of intrinsic localization motifs and the spatial information provided by the fused domains. In support, while both PKCθ V3 and PLCδPH did not display central localization in isolation, they could enhance central localization of LAT and SLP-76 (Figs. 4, 6). Similarly, artificially enhanced cSMAC formation through expression of LAT V3 did not lead to the recruitment of signaling intermediates with intrinsically weak cSMAC preference such as Vav1 (Fig. 4C). Our inability to force cSMAC localization implies that even upon addition of new protein interaction domains the overall molecular composition of supramolecular complexes in the cSMAC is fairly well conserved: A core of protein-protein interactions may be required for complex formation such that addition of new protein interactions can only enhance complex stability but not generate complexes of fundamentally different composition. Supramolecular assemblies in the cSMAC thus likely exist in a delicate balance between compositional conservation for complex identity and some redundancy in protein interaction motifs and stoichiometry to allow flexible regulation of stability.

Reconstitution of central LAT and SLP-76 clustering under attenuated T cell activation conditions could restore IL-2 mRNA generation. The association of lack of Grb2 centrality with lack of an effect on IL-2 generation provides a specificity control. Protein clustering in the cSMAC thus was a critical component of efficient T cell signaling. For the cSMAC to enhance T cell function experimental reconstitution of cSMAC formation needed to closely mimic cSMAC features observed with non-targeted constructs under full stimulus conditions. Time-dependent roles of the cSMAC have been proposed before (Freiberg et al., 2002) yet are difficult to compare to the work here as the molecules investigated differed. In our work under full stimulus conditions LAT-GFP and SLP-76-GFP were efficiently recruited to the interface center within the first two minutes of T cell activation with diminished recruitment thereafter. Recruitment of LAT and SLP-76 to the interface center upon attenuation of T cell activation by fusion with the Vav1 SH3SH2SH3 and PLCδ PH domains closely reproduced these dynamics (Figs. 4B, 6) and largely restored IL-2 mRNA generation (Figs. 5A, 6). However, recruitment of LAT to the interface center to a greater extent and duration by fusion with PKCθ V3 (Fig. 4B) could not enhance IL-2 mRNA generation (Fig. 5A). We therefore suggest that the cSMAC displays two different time-dependent roles. Within the first one to two minutes of T cell activation it efficiently brings together the large number of proximal T cell signaling intermediates required for efficient T cell activation. Subsequently, a substantial number of key signalling intermediates including SLP-76, Itk, PLC*γ*, and Vav1 leave the cSMAC and move to smaller signalling complexes supported by an interface-wide lamellal actin network (Roybal et al., 2015). Retention of a more limited subset of signalling intermediates in the cSMAC after this time thus may render them less accessible to their interaction partners and therefore diminish sustained signal transduction. Signal enhancing and attenuating roles of the cSMAC thus may both occur as regulated by the specific time-dependent composition of the complex.

Itk and costimulation likely contribute to cSMAC formation by overlapping yet partially distinct means. Recruitment to the interface center peaks within the first two minutes of cell coupling for both ligand-engaged CD28 (Purtic et al., 2005; Singleton et al., 2009) and full length Itk (Roybal et al., 2015). Both thus can be expected to provide protein-protein interactions during the early signal amplifying stage of the cSMAC. Itk also has enzymatic activity to directly modify cSMAC components. Both costimulation and Itk regulate actin dynamics. However, while costimulation controls core actin turnover through the Arp2/3 complex and Cofilin (Roybal et al., 2016), Itk only regulates a SLAT-dependent subset of actin dynamics (Singleton et al., 2011). CD28 thus can be expected to contribute more strongly to cSMAC directed transport in complex assembly (Roybal et al., 2016)(Fig. 2C).

In summary, our work establishes that the cSMAC formed in the activation of T cells by APCs consists of multiple supramolecular complexes driven by extensive membrane undulations with a dynamically changing composition. Compositionally rich complexes in the first two minutes of cell coupling enhanced T cell activation by facilitating efficient signaling interactions whereas thereafter compositionally poorer complexes may sequester signaling intermediates.

## Acknowledgements

We acknowledge the University of Bristol FACS and Wolfson BioImaging facilities for providing equipment and technical support. The work was supported by grants from the US National Science Foundation (MCB1121793 to CW, MCB1121919 and MCB1616492 to RFM), ERC (CW: PCIG11-GA-2012-321554), BBSRC (CW: BB/P011578/1) and US National Institutes of Health (P41 GM103712).

## Author contributions

DJC, LEM, RFM, CW conceived the study and designed experiments, DJC, LEM, SLT, GB, CM, HT, LM, XR, KLS, MD, CW performed experiments and analyzed data, AJH executed statistical analysis, PLS provided Itk knock out mice, CW wrote the manuscript.

## Methods

### Antibodies and reagents

Antibody used for quantitative Western blotting were α-LAT pY191 (Cell Signaling, #3584, RRID:AB_2157728), α-GAPDH Clone 14C10 (Cell Signaling, #2118, RRID:AB_561053), and α-alpha Tubulin Clone DM1A (ThermoFisher Scientific, #62204, RRID:AB_1965960). Antibodies used for the blockade of B7-1- and B7-2-dependent CD28 costimulation were anti-mouse CD80 Clone 16-10-A1 (BD Pharmingen #553736) and anti-CD86 Clone GL1 (BD Pharmingen #553689). Protein transduction versions of constitutively active cofilin (S3A) and Rac1 (Q61L) were purified from *E. coli* and introduced into primary 5C.C7 T cells by 30 min incubation as previously described (Roybal et al., 2016).

### Mice and cells

Itk-deficient 5C.C7 mice were generated by crossing B10.BR 5C.C7 TCR transgenic mice (Seder et al., 1992)(RRID:MGI:3799371) with Itk-deficient B6 mice (Schaeffer et al., 1999)(RRID:MGI:4356470). T cells expanded from the lymph nodes of wild type or Itk-deficient 5C.C7 TCR transgenic mice were used for all experiments. The 5C.C7 TCR recognizes the moth cytochrome c peptide fragment (amino acid residues 88 to 103, ANERADLIAYLKQATK) in the context of I-E^k^. Single-cell suspensions were made from the lymph nodes of 6- to 8-week-old mice of either gender. The cells were adjusted to 4 × 10^6^ cells/ml and MCC peptide was added to a final concentration of 3 µM. T cells were transduced with MMLV-derived retroviruses for the expression of signaling sensors, commonly signaling intermediates fused with GFP as described in detail (Ambler et al., 2017; Roybal et al., 2015; Singleton et al., 2009). All animals were maintained in pathogen-free animal facilities at the University of Bristol under a University mouse breeding Home Office License. The CH27 B cell lymphoma cell line (RRID:CVCL_7178) was used in all experiments as APCs. To load the APCs, the cells were incubated in the presence of 10 µM MCC peptide for at least 4 hours. All cells were maintained in medium composed of RPMI with L-glutamine, 10% fetal bovine serum (FBS, Hyclone), penicillin (100 IU/mL), streptomycin (100 µg/ml), and 0.5 µM *β*-mercaptoethanol. Interleukin-2 (IL-2)(TECIN recombinant human IL-2, NCI Biological Resource Branch) was added at a final concentration of 0.05 U/ml during parts of the retroviral transduction procedure.

### Time-lapsed imaging of T cell:APC interactions

Our imaging and image analysis protocols have recently been described in great detail in a dedicated publication (Ambler et al., 2017). Briefly, time-lapse fluorescence microscopy was performed with retrovirally transduced T cells, FACS-sorted to the lowest detectable sensor expression of 2 µM, and CH27 cells loaded with 10 µM MCC. The T cells and CH27 cells were imaged in imaging buffer (PBS, 10% FBS, 1 mM CaCl_2_, 0.5 mM MgCl_2_) on 384-well glass-bottom plates. All imaging was performed on a Perkin Elmer UltraVIEW ERS 6FE spinning disk confocal systems fitted onto a Leica DM I6000 microscope body equipped with full environmental control and a Hamamatsu C9100-50 EMCCD. A Leica 40x HCX PL APO oil objective (NA = 1.25) was used for all imaging. Automated control of the microscope was performed with Volocity software (Perkin Elmer). For experiments in which the B7-1- and B7-2-dependent activation of CD28 was blocked, peptide-loaded CH27 cells were incubated on ice for 30 min in the presence of anti- CD80 Clone 16-10-A1 (10 µg/ml) and anti- CD86 Clone GL1 (10 µg/ml)(BD Pharmingen) antibody before the CH27 cells were transferred to the imaging plate with the T cells. For experiments in which cells were reconstituted with protein transduction versions of constitutively active Rac and cofilin, T cells were incubated for 30 min at 37°C with the protein transduction reagents at the indicated concentrations in the imaging plate before the addition of the peptide-loaded CH27 cells. Each time-lapse image sequence was generated by taking a differential interference contrast brightfield image and a 3D image stack of the GFP channel every 20 s for 46 frames at 37°C. Voxels in these 3D images were of size 0.34 µm in the horizontal plane and 1 µm along the optical axis.

### Image Analysis

The location and frame number of each T cell:APC couple were identified as when either the T cell:APC interface had reached its full width or the cells had been in contact for 40 s, whichever came first. Patterns of signaling sensor enrichment were assessed according to previously established quantitative criteria (Fig. 2 in (Singleton et al., 2009)) as depicted in the Fig. S1A. Briefly, the six, mutually exclusive interface patterns were: accumulation at the center of the T cell-APC interface (central), accumulation in a large T cell invagination (invagination), accumulation that covered the cell cortex across central and peripheral regions (diffuse), accumulation in a broad interface lamella (lamellal), accumulation at the periphery of the interface (peripheral) or in smaller protrusions (asymmetric). Briefly, for each cell couple at each time point we first determined whether fluorescence intensity in the area of accumulation was >40% above the cellular fluorescence background. If so, the geometrical features of the area of accumulation, fraction of the interface covered, location within the interface, and extension of the area of accumulation away from the interface (Fig. 2 in (Singleton et al., 2009)), were used to assign the cell couple to one of the mutually exclusive patterns. Systems-scale cluster analysis was performed with Cluster (Michael Eisen, UC Berkeley) as established (Singleton et al., 2009). To measure interface enrichment of LAT and spatially targeted versions thereof we used a recently developed computational image analysis routine (Roybal et al., 2016). Very briefly, starting with the manual cell couple identification described above T cells were segmented, reoriented with the T cell:APC interface facing up using the ‘two-point synapse annotation’ procedure (Roybal et al., 2016), and the cell shape was standardized to a half spheroid to allow voxel-by-voxel comparison across all cell couples analysed. After transformation to the standard shape, the fluorescence distribution in each cell at a given time point was represented as a standardized vector (of length 6628) formed from the intensity values of each of the voxels within the template shape, where the intensities for each time point were normalized so that the values of the vector were probabilities (that is, fractions of total intensity). To measure interface enrichment, we defined an interface enrichment region as the 10% most fluorescent voxels of the average probability distribution across all cells, for all time points, and for all sensors. We defined enrichment to be the ratio of the mean probability in the distribution of that sensor for that cell at that time point within the interface enrichment region and the mean probability in the entire cell.

### STED imaging

CH27 APCs were adhered to α-CD19 antibody-coated coverslips. T cells were then allowed to interact with APCs for 4.5 minutes and fixed with 4% PFA for 20 minutes at 4 °C followed by PFA quenching using ammonium chloride for 10 minutes at 4 °C. Cells were permeabilized for 20 min in 0.02% Triton X-100 in PBS at 4 °C. T cells were blocked in 1% BSA in PBS for 30 min at room temperature and probed with primary antibodies against LAT (1:100, Cell Signalling #9166, RRID:AB_2283298) or phospho-LAT Tyr 191 (1:50, Cell Signalling #3584, RRID:AB_2157728) diluted in 1% BSA with Fc block (Rat Anti-Mouse CD16/CD32, #553141, BD Bioscience, RRID:AB_394656) at the same dilution in PBS for overnight at 4 °C. Cells were washed with PBS three times and incubated with secondary antibody, Donkey anti-rabbit IgG, Alexa Fluor 488 (Molecular Probes #R37118, 1:1000, RRID:AB_2556546) in 1% BSA with Fc block (1:500) for 1 hour at room temperature. Coverslips were washed with PBS before mounting using ProLong Gold (Thermo Fisher) and cured for 24 hours at room temperature.

Fixed cells were imaged through a 100x HC PL APO CS2 1.4 NA objective on a Leica SP8 AOBS confocal laser scanning microscope. Alexa Fluor 488 was excited using a white light laser with an emission filter between 498 - 520 nm and STED depletion was achieved using a 592 nm continuous wave fibre laser. Images were first de-convolved using Hyugen Professional followed by automated puncta analysis with ImageJ (NIH). LAT puncta were detected and measured using Wolfson Bioimaging ImageJ plugins (Modular Image Analysis). Individual puncta were identified using the Otsu algorithm (Otsu, 1979) with a threshold multiplier of 3.5 A.U. followed by a filtration mode of the Watershed 3D method to identify separate puncta. Small puncta detected in the APCs that don’t express LAT were used to derive a detection size threshold for LAT complexes such that all puncta smaller than the 95 percentile of the APC puncta size distribution were excluded from the analysis. Thus, complexes smaller than 0.04/0.06µm^3^ in the LAT/pLAT data were excluded from the analysis. Repeating the analysis with a 99-percentile cut-off didn’t change the conclusions reached.

### CLEM

The interaction of 5C.C7 T cells with CH27 B cell APCs was imaged live by spinning disk confocal microscopy as described above using a 35 mm glass bottom finder dish (Mattek). Upon formation of a cSMAC, cells were immediately fixed using 2.5% Glutaraldehyde (Agar Scientific) in 0.1M cacodylate buffer, stained in 1% osmium tetroxide (EMS) in cacodylate buffer, dehydrated and embedded in Epon812 resin (TAAB) for 24 hours at 60°C. Samples were removed from EPON and trimmed to section of interest. Trimmed samples were sectioned at 300 nm using an Ultramicrotome (Leica, EM UC7) with a diamond knife (DiATOME) and stained with uranyl acetate and lead citrate (Agar Scientific). Sections were analysed using a FEI Tecnai 12 120 kV BioTwin equipped with a bottom-mount 4*4 K Eagle CCD camera. The tomogram data series was acquired using a FRI Tecnai 20 TEM between −50° to + 50° with a 2.0° increment. The data was reconstructed using IMOD etomo software.

Segmentation was made using AMIRA software (VSG), while for visualization, a combination of IMOD, AMIRA and Image J were used.

### IL-2 ELISA

Live wild type or Itk-deficient 5C.C7 T cells were FACS sorted to generate comparable cell numbers across each assay. CH27 B cells were peptide loaded with 10μM MCC for four hours or overnight. T and B cells were mixed in round bottom 96-well plates at 1 × 10^4^ T cells to 5 × 10^4^ B cells. For costimulation blockade 10μg/ml α-CD80 and α-CD86 were added to each well. The cells were then incubated for 18 hours and IL-2 amounts in the supernatant were determined using a mouse IL-2 OptEIA ELISA kit from BD Biosciences as per manufacturer’s instructions.

### IL-2 mRNA determination

CH27 B cell lymphoma APCs were peptide loaded overnight with 10μM MCC peptide. Live wild type or Itk-deficient 5C.C7 T cells, non-transduced or expressing adaptor protein-GFP or targeted variants thereof, were FACS sorted to generate comparable cell numbers across each assay. 1 × 10^4^ T cells and 5 × 10^4^ APCs cells were centrifuged for 30 seconds at 1,000 rpm to maximize cell-to-cell contact and incubated at 37°C for 2 hours. For costimulation blockade 10μg/ml α-CD80 and α-CD86 were added to each well. mRNA was isolated using the Qiagen RNeasy Micro Kit (Qiagen, UK) according to manufacturer’s instructions. cDNA was generated using an Invitrogen AMV First-Strand cDNA synthesis kit (Life Technologies, UK) according to manufacturer’s instructions. IL-2 mRNA amounts were determined with a SYBR Green PCR master mix from Life Technologies (4344463) relative to mRNA for β-2 microglobulin on a DNA Engine Opticon II System (Bio-Rad) using the following oligonucleotides, IL-2: AGCTGTTGATGGACCTA and CGC AGA GGT CCA AGT TCA T, β-2 microglobulin: GCTATCCAGAAAACCCCTCAA and CGG GTG GAA CTG TGT GTT ACG T.

### Western blotting analysis

CH27 B cell lymphoma APCs were peptide loaded overnight with 10μM MCC peptide. Live wild type or Itk-deficient 5C.C7 T cells, non-transduced or expressing LAT-GFP or a targeted variant thereof, were FACS sorted to generate comparable cell numbers across each assay. 1 × 10^6^ T cells and 1 × 10^6^ APCs cells were centrifuged for 30 seconds at 1,000 rpm to maximize cell-to-cell contact and incubated at 37°C for the indicated time. Subsequently, samples were immediately lysed with cold RIPA lysis buffer (Millipore) plus protease/ phosphatase inhibitor cocktail (Cell Signaling) for 30 minutes on ice. To remove the insoluble fraction, samples were centrifuged at 20,000 g for 15 minutes. Supernatant were run on SDS/PAGE gels, transferred to PDVF membranes and blotted according to standard protocols. Blots were stripped and reprobed with an anti-GAPDH or anti-α tubulin antibody to normalize for sample loading.

### Statistics

The frequency of occurrence of interface accumulation patterns was analyzed pairwise with a proportion’s z-test as reported in the supplementary tables. p values were not corrected for multiple comparisons as the corresponding pFDR q-values (Storey, 2004) were similar. IL-2 mRNA amounts were first logarithmically transformed to stabilize the variance and approximate to the normal distribution and then analyzed by 1-way ANOVA with Tukey’s adjustment for multiple comparisons or 2-way ANOVA with the Sidak adjustment for multiple comparisons depending on the number of variables compared. IL-2 ELISA data were first logarithmically transformed and then analyzed by 1-way ANOVA with Tukey’s adjustment for multiple comparisons. LAT phosphorylation data were first logarithmically transformed and then analyzed by 1-way ANOVA with Tukey’s adjustment for multiple comparisons separately for each time point.

### Data availability statement

All imaging data are openly accessible through http://murphylab.cbd.cmu.edu/data/TcellLAT2018/ IL-2 and LAT phosphorylation data that support the findings of this study are available from the corresponding author upon reasonable request.

---

**Table S1 Sensors used in Fig. 1C, D**. Publications describing the sensors used in Fig. 1C, D and representative 5C.C7 T cell imaging data are given.

**Table S2 Statistical significance** of differences in LAT accumulation under different T cell activation conditions are given for the indicated patterns as determined by proportion’s z-test. No entry indicates p > 0.05.

**Table S3 Statistical significance** of differences in accumulation of spatially targeted as compared to non-targeted LAT under different T cell activation conditions are given for the indicated patterns as determined by proportion’s z-test. No entry indicates p > 0.05.

**Table S4 Statistical significance** of differences in accumulation of Grb2, Lck and Vav1 in the presence as compared to absence of LATV3 under different T cell activation conditions are given for the indicated patterns as determined by proportion’s z-test. No entry indicates p > 0.05.

**Table S5 Statistical significance** of differences in SLP-76 accumulation under different T cell activation conditions are given for the indicated patterns as determined by proportion’s z-test on top. Statistical significance of differences in accumulation of spatially targeted as compared to non-targeted SLP-76 under different T cell activation conditions are given for the indicated patterns as determined by proportion’s z-test on the bottom. No entry indicates p > 0.05.

**Table S6 Statistical significance** of differences in Grb2 accumulation under different T cell activation conditions are given for the indicated patterns as determined by proportion’s z-test on top. Statistical significance of differences in accumulation of spatially targeted as compared to non-targeted Grb2 under different T cell activation conditions are given for the indicated patterns as determined by proportion’s z-test on the bottom. No entry indicates p > 0.05.

**Fig. S1 Sensor accumulation at the T cell:APC interface under full stimulus conditions. A** The panel graphically represents the six categories used to classify spatiotemporal sensor distribution as underpinned by defined cell biological structures at the T cell:APC interface (Roybal et al., 2013; Singleton et al., 2009). The antigen-presenting cell above the T cell is not shown. Central reflects the cSMAC, lamellal an F-actin-based lamella extending from the undulating T cell plasma membrane deep into the T cell, peripheral the part of the actin network stabilizing the interface edge. Diffuse reflects cortical accumulation, invagination enrichment in a transient large T cell invagination and asymmetric individual small lamellae. For all main figures the patterns ‘diffuse’ and ‘lamellal’ are combined as ‘diffuse/lamellal’, the patterns ‘asymmetric’ and ‘peripheral’ as ‘periphery’. **B** The graphs display the percentage of cell couples that displayed accumulation of the indicated sensor (Table S1) with the indicated patterns (A) relative to tight cell couple formation for wild type 5C.C7 T cells activated with CH27 B cell APCs (10 µM MCC peptide). 49-141 cell couples were analyzed per condition, 869 total.

**Fig. S2 Sensor accumulation at the T cell:APC interface under costimulation blocked conditions.** The graphs display the percentage of cell couples that displayed accumulation of the indicated sensor (Table S1) with the indicated patterns (Fig. S1A) relative to tight cell couple formation for wild type 5C.C7 T cells activated with CH27 B cell APCs (10 µM MCC peptide) in the presence of 10 µg/ml anti-CD80 plus anti-CD86. 39-107 cell couples were analyzed per condition, 869 total.

**Fig. S3 Sensor accumulation at the T cell:APC interface in the absence of Itk.** The graphs display the percentage of cell couples that displayed accumulation of the indicated sensor (Table S1) with the indicated patterns (Fig. S1A) relative to tight cell couple formation for Itk-deficient 5C.C7 T cells activated with CH27 B cell APCs (10 µM MCC peptide). 30-65 cell couples were analyzed per condition, 591 total.

**Fig. S4 Representative phospho-LAT Y191 Western blot.** Wild type or Itk-deficient 5C.C7 T cells were activated with CH27 B cell and 10 µm MCC in the presence or absence of 10 µg/ml anti-CD80 plus anti-CD86. At the given time points (in minutes) T cell:APC cell extracts were blotted for LAT phosphorylation at Y191. Blots were stripped and reblotted with an anti-alpha tubulin antibody as a loading control. Phospho-LAT Y191 band intensities as normalized to loading control are given under the phospho-LAT blots. Molecular weights of marker bands (in kD) are given at the right. One representative blot of eight is shown.

**Fig. S5 Patterning of isolated targeting domains and interface recruitment of LAT-GFP and spatially targeted version thereof. A** The graphs give the percentage of cell couples that displayed accumulation of the isolated targeting domains, LAT V3, LAT Vav, and LAT PLCδPH as indicated, with the indicated patterns as in Fig. 2B relative to tight cell couple formation for wild type 5C.C7 T cells activated with CH27 B cell APCs (10 µM MCC peptide). 30-57 cell couples were analyzed per condition, 133 total. **B, C** For the quantification of the accumulation of LAT and targeted versions thereof at the T cell:APC interface we applied a computational image quantification as recently described in detail (Roybal et al., 2016). We identified a core region (B) of sensor enrichment as defined as the 10% most fluorescent voxels of the average probability distribution across all cells, for all time points, and for all sensors. Using this core region, we calculated the ratio of the amount of the sensor in the region to the average amount across the whole cell. This was done for all time points either under conditions of full stimulus or costimulation blockade as indicated. Imaging data analyzed are the same as in Figs. 2 and 3. **D** Wild type 5C.C7 T cells retrovirally transduced to express the indicated LAT constructs were activated with CH27 B cell APCs and 10 µm MCC in the presence of 10 µg/ml anti-CD80 plus anti-CD86. At the given time points T cell:APC cell extracts were blotted for LAT phosphorylation at Y191. Blots were stripped and reblotted with an anti-alpha tubulin antibody as a loading control. One representative blot of two is shown.

**Fig. S6 IL2 mRNA amounts upon expression of the isolated targeting domains.** Wild type or Itk-deficient (‘Itk ko’) 5C.C7 T cells were transduced to express the isolated targeting domains, LAT V3, LAT Vav, and LAT PLCδPH as indicated, and reactivated by CH27 APCs (10 µM MCC peptide) in the absence or presence of 10 µg/ml anti-CD80 plus anti-CD86 (‘costimulation blockade’). IL-2 mRNA amounts are given relative to IL-2 mRNA in non-transduced 5C.C7 T cells under full stimulus conditions. 3-6 experiments were averaged per condition. None of the differences were significant as calculated with 2-way ANOVA.

**Video S1.** Representative interactions of 5C.C7 T cells retrovirally transduced to express the indicated GFP fusion proteins with CH27 B cell lymphoma APCs and 10 μM MCC peptide in the presence or absence of 10 µg/ml anti-CD80 plus anti-CD86 (‘costimulation blockade’) are shown in videos S1 and S4 to S10. DIC images are shown on the top, with matching top-down, maximum projections of 3D sensor fluorescence data on the bottom. The sensor fluorescence intensity is displayed in a rainbow-like, false-color scale (increasing from blue to red). 20 s intervals in video acquisition are played back as 2 frames per second. The 5C.C7 T cell in Video S1 is transduced with LAT-GFP and responds to a full stimulus. Cell coupling occurs in frame 4 (2s indicated video time).

**Video S2.** Representative EM tomograms are shown in videos S2 and S3. Reconstructed Z-sections are first shown without tracing and subsequently with the T cell and APC plasma membranes at the interface traced in blue and red, respectively. The 5C.C7 T cell in Video S2 is transduced with LAT-GFP and responds to a full stimulus. It displayed central LAT-GFP accumulation at the time of fixation.

**Video S3.** The video is displayed similar to Video S2. The 5C.C7 T cell in Video S3 is transduced with LAT V3-GFP and responds to a full stimulus upon costimulation blockade. It displayed central LAT-GFP accumulation at the time of fixation.

**Video S4.** The video is displayed similar to Video S1. The 5C.C7 T cell in Video S4 is transduced with LAT V3-GFP and responds to a full stimulus upon costimulation blockade. Cell coupling occurs in frame 4 (2s indicated video time).

**Video S5.** The video is displayed similar to Video S1. The 5C.C7 T cell in Video S5 is transduced with LAT Vav-GFP and responds to a full stimulus upon costimulation blockade. Cell coupling occurs in frame 5 (2s indicated video time).

**Video S6.** The video is displayed similar to Video S1. The 5C.C7 T cell in Video S6 is transduced with LAT PLCδ PH-GFP and responds to a full stimulus upon costimulation blockade. Cell coupling occurs in frame 5 (2s indicated video time).

**Video S7.** The video is displayed similar to Video S1. The 5C.C7 T cell in Video S7 is transduced with SLP-76-GFP and responds to a full stimulus. Cell coupling occurs in frame 4 (2s indicated video time).

**Video S8.** The video is displayed similar to Video S1. The 5C.C7 T cell in Video S8 is transduced with SLP-76 V3-GFP and responds to a full stimulus upon costimulation blockade. Cell coupling occurs in frame 4 (2s indicated video time).

**Video S9.** The video is displayed similar to Video S1. The 5C.C7 T cell in Video S9 is transduced with SLP-76 Vav-GFP and responds to a full stimulus upon costimulation blockade. Cell coupling occurs in frame 5 (2s indicated video time).

**Video S10.** The video is displayed similar to Video S1. The 5C.C7 T cell in Video S10 is transduced with Grb2-GFP and responds to a full stimulus. Cell coupling occurs in frame 5 (2s indicated video time).

**Video S11.** The video is displayed similar to Video S1. The 5C.C7 T cell in Video S11 is transduced with Grb2 V3-GFP and responds to a full stimulus upon costimulation blockade. Cell coupling occurs in frame 4 (2s indicated video time).

**Video S12.** The video is displayed similar to Video S1. The 5C.C7 T cell in Video S12 is transduced with Grb2 Vav-GFP and responds to a full stimulus upon costimulation blockade. Cell coupling occurs in frame 5 (2s indicated video time). A second T cell activates at the left edge of the filed in frame 7 (3s indicated video time).

