## Supplementary tables for "Transient protein accumulation at the center of the T cell antigen presenting cell interface drives efficient IL-2 secretion"

#### Table S1 – to Fig. 1

| Signaling intermediate | Sensor | Location of classification data/representative video |
|---|---|---|
| **Actin** | GFP-Actin | J. Immunol. (2003) *171,* 2287-95 |
| **Akt** | Akt-GFP | PLoS One (2015) *10,* e0133299 |
| **Arp3** | Arp3-GFP | Sci. Signal. (2016) *9*, rs3 |
| **Capping protein 1 α** | Capping protein 1 α-GFP | Sci. Signal. (2016) *9*, rs3 |
| **CD2** | CD48-GFP | Sci. Signal. (2009) *2*, ra15 |
| **Cofilin** | Cofilin-GFP | Sci. Signal. (2016) *9*, rs3 |
| **Coronin 1A** | Coronin 1A-GFP | Sci. Signal. (2016) *9*, rs3 |
| **Ezrin** | Ezrin-GFP | PLoS One (2015) *10,* e0133299 |
| **Grb2** | Grb2-GFP | PLoS One (2015) *10,* e0133299 |
| **HS1** | HS1-GFP | Sci. Signal. (2016) *9*, rs3 |
| **Itk** | Itk-GFP | PLoS One (2015) *10,* e0133299 |
| **LAT** | LAT-GFP | Sci. Signal. (2009) *2*, ra15 |
| **Lck** | Lck-GFP | PLoS One (2015) *10,* e0133299 |
| **Myosin II RLC** | Myosin II RLC-GFP | Sci. Signal. (2016) *9*, rs3 |
| **NFκB p65** | GFP-p65 | PLoS One (2015) *10,* e0133299 |
| **$PIP_2$** | GFP-PLCδ-PH | PLoS One (2015) *10,* e0133299 |
| **PKC Θ** | PKC Θ-GFP | Sci. Signal. (2009) *2*, ra15 |
| **SLP-76** | SLP-76-GFP | PLoS One (2015) *10,* e0133299 |
| **TCR as TCRζ** | TCRζ-GFP | Sci. Signal. (2009) *2*, ra15 |
| **Vav1** | Vav1-GFP | PLoS One (2015) *10,* e0133299 |
| **WASP** | GFP-WASP | Sci. Signal. (2016) *9*, rs3 |
| **WAVE-2** | GFP-WAVE2 | Sci. Signal. (2016) *9*, rs3 |

### Table S2 – to Fig. 2

| Condition | Comparison | Pattern | -40 | -20 | 0 | 20 | 40 | 60 | 80 | 100 | 120 | 180 | 300 | 420 |
|---|---|---|---|---|---|---|---|---|---|---|---|---|---|---|
| LAT, anti-B7 | LAT, full stim. | any | | | 0.000 | 0.000 | 0.000 | 0.000 | 0.000 | 0.000 | 0.000 | 0.005 | 0.04 | 0.02 |
| | | central | | | 0.000 | 0.000 | 0.001 | 0.006 | | | | | | |
| LAT, Itk ko, full stim. | LAT, full stim. | any | | | 0.000 | | | | | | | | | |
| | | central | | | 0.005 | 0.02 | | | | | | | | |
| LAT, Itk ko, anti-B7 | LAT, full stim. | any | | | 0.000 | 0.000 | 0.000 | 0.000 | 0.000 | 0.000 | 0.000 | 0.000 | 0.02 | 0.005 |
| | | central | | | 0.000 | 0.000 | 0.05 | 0.01 | 0.03 | 0.05 | | 0.03 | | |
| LAT, Itk ko, anti-B7, Rac Cofilin | LAT, full stim. | any | | | 0.001 | 0.001 | | | | | | 0.02 | 0.007 | 0.008 |
| | | central | | | 0.008 | | | | 0.02 | 0.000 | 0.000 | 0.01 | 0.05 | |
| LAT, Itk ko, anti-B7, Rac Cofilin | LAT, Itk ko, anti-B7 | any | | | | | 0.006 | 0.000 | 0.000 | 0.000 | 0.000 | 0.000 | 0.000 | 0.000 |
| | | central | | | | 0.03 | 0.01 | 0.002 | 0.000 | 0.000 | 0.000 | 0.000 | 0.02 | |

### Table S3 – to Fig. 4

#### LAT V3

| Condition | Comparison | Pattern | -40 | -20 | 0 | 20 | 40 | 60 | 80 | 100 | 120 | 180 | 300 | 420 |
|---|---|---|---|---|---|---|---|---|---|---|---|---|---|---|
| **Restoration to LAT full stimulus** | | | | | | | | | | | | | | |
| LAT V3, full stimulus | LAT, full stimulus | any | | | 0.03 | 0.001 | 0.000 | 0.000 | 0.000 | 0.000 | 0.000 | 0.000 | 0.000 | 0.000 |
| | | central | | | | | 0.004 | 0.001 | 0.000 | 0.000 | 0.000 | 0.000 | 0.03 | 0.002 |
| LAT V3, Itk ko, full stimulus | LAT, full stimulus | any | | 0.002 | | | 0.04 | 0.01 | 0.004 | 0.003 | 0.002 | 0.001 | 0.001 | 0.000 |
| | | central | | | 0.01 | | 0.007 | 0.006 | 0.000 | 0.000 | 0.000 | 0.000 | 0.004 | 0.006 |
| LAT V3, anti-B7 | LAT, full stimulus | any | | 0.006 | | 0.008 | 0.002 | 0.004 | 0.003 | 0.000 | 0.000 | 0.000 | 0.006 | 0.006 |
| | | central | | | | | 0.04 | 0.03 | 0.02 | 0.000 | 0.000 | 0.000 | | |
| LAT V3, Itk ko anti-B7 | LAT, full stimulus | any | | | 0.000 | 0.006 | | | | | | 0.01 | 0.009 | 0.02 |
| | | central | | | 0.000 | | 0.000 | 0.000 | 0.000 | 0.000 | 0.000 | 0.000 | 0.03 | 0.003 |
| **Enhancement under matched stimuli** | | | | | | | | | | | | | | |
| LAT V3, Itk ko, full stimulus | LAT, Itk ko, full stimulus | any | | 0.004 | 0.007 | | | | 0.03 | 0.01 | 0.004 | 0.04 | 0.01 | 0.001 |
| | | central | | | | 0.03 | 0.01 | 0.005 | 0.001 | 0.000 | 0.001 | 0.001 | 0.006 | 0.004 |
| LAT V3, anti-B7 | LAT, anti-B7 | any | | | | | 0.002 | 0.001 | 0.000 | 0.000 | 0.000 | 0.000 | 0.006 | 0.003 |
| | | central | | | | 0.02 | 0.000 | 0.000 | 0.000 | 0.000 | 0.000 | 0.000 | 0.003 | 0.004 |
| LAT V3, Itk ko, anti-B7 | LAT, Itk ko, anti-B7 | any | | 0.000 | 0.000 | 0.000 | 0.000 | 0.000 | 0.000 | 0.000 | 0.000 | 0.000 | 0.001 | 0.000 |
| | | central | | | 0.04 | 0.001 | 0.000 | 0.000 | 0.000 | 0.000 | 0.000 | 0.000 | 0.01 | 0.000 |

#### LAT Vav

| Condition | Comparison | Pattern | -40 | -20 | 0 | 20 | 40 | 60 | 80 | 100 | 120 | 180 | 300 | 420 |
|---|---|---|---|---|---|---|---|---|---|---|---|---|---|---|
| **Restoration to LAT full stimulus** | | | | | | | | | | | | | | |
| LAT Vav, full stimulus | LAT, full stimulus | any | | | 0.002 | 0.03 | 0.004 | 0.002 | 0.001 | 0.003 | 0.000 | 0.002 | | |
| | | central | | | 0.05 | 0.01 | | | | | | | | |
| LAT Vav, Itk ko, full stimulus | LAT, full stimulus | any | | | 0.000 | 0.001 | | | | | | | 0.008 | |
| | | central | | | 0.005 | | | 0.001 | 0.001 | 0.000 | 0.000 | 0.001 | | |
| LAT Vav, anti-B7 | LAT, full stimulus | any | | | | | | | | | | | | |
| | | central | | | | | 0.002 | 0.002 | 0.002 | 0.000 | 0.002 | | | |
| LAT Vav, Itk ko anti-B7 | LAT, full stimulus | any | | | 0.000 | 0.000 | | | | | | 0.04 | | |
| | | central | | | 0.000 | | 0.02 | 0.006 | 0.000 | 0.000 | 0.000 | 0.002 | | 0.05 |
| **Enhancement under matched stimuli** | | | | | | | | | | | | | | |
| LAT Vav, Itk ko, full stimulus | LAT, Itk ko, full stimulus | any | | | | 0.01 | 0.05 | | | | | | 0.05 | |
| | | central | | | | | | 0.001 | 0.05 | 0.02 | 0.03 | 0.005 | | |
| LAT Vav, anti-B7 | LAT, anti-B7 | any | | | 0.001 | 0.01 | 0.000 | 0.000 | 0.001 | 0.001 | 0.005 | | | 0.03 |
| | | central | | | 0.000 | 0.000 | 0.000 | 0.000 | 0.001 | 0.000 | 0.000 | | | |
| LAT Vav, Itk ko, anti-B7 | LAT, Itk ko, anti-B7 | any | | | | 0.03 | 0.001 | 0.000 | 0.000 | 0.000 | 0.000 | 0.000 | 0.001 | 0.000 |
| | | central | | | | 0.005 | 0.000 | 0.000 | 0.000 | 0.000 | 0.000 | 0.000 | 0.03 | 0.007 |

#### LAT PLCdPH

| Condition | Comparison | Pattern | -40 | -20 | 0 | 20 | 40 | 60 | 80 | 100 | 120 | 180 | 300 | 420 |
|---|---|---|---|---|---|---|---|---|---|---|---|---|---|---|
| **Restoration to LAT full stimulus** | | | | | | | | | | | | | | |
| LAT PLCd, full stimulus | LAT, full stimulus | any | | | 0.000 | 0.000 | 0.000 | 0.002 | 0.03 | 0.03 | | | | |
| | | central | | | 0.000 | 0.05 | | | | | | | | |
| LAT PLCd, Itk ko, full stimulus | LAT, full stimulus | any | | | 0.000 | 0.01 | | | 0.001 | | 0.03 | | | |
| | | central | | | 0.001 | | | | 0.05 | 0.04 | | | | |
| LAT PLCd, anti-B7 | LAT, full stimulus | any | | | | 0.03 | 0.01 | 0.002 | 0.000 | 0.000 | 0.000 | 0.000 | 0.001 | 0.05 |
| | | central | | | | 0.009 | | | | | | | | 0.04 |
| LAT PLCd, Itk ko anti-B7 | LAT, full stimulus | any | | | 0.000 | 0.009 | | | | | | 0.005 | | |
| | | central | | | 0.001 | 0.02 | | | | | | | | |
| **Enhancement under matched stimuli** | | | | | | | | | | | | | | |
| LAT PLCd, Itk ko, full stimulus | LAT, Itk ko, full stimulus | any | | | | | | | 0.005 | | 0.05 | | | |
| | | central | 0.05 | | | | | | | | | | | |
| LAT PLCd, anti-B7 | LAT, anti-B7 | any | | | 0.009 | | | 0.03 | | | | | | |
| | | central | | | 0.01 | | | | | | | | | |
| LAT PLCd, Itk ko anti-B7 | LAT, Itk ko, anti-B7 | any | | | | 0.005 | 0.002 | 0.000 | 0.000 | 0.000 | 0.000 | 0.000 | 0.002 | 0.002 |
| | | central | | | | | | | | 0.003 | 0.05 | | | |

### Table S4 – to Fig. 5

| Condition | Comparison | Pattern | -40 | -20 | 0 | 20 | 40 | 60 | 80 | 100 | 120 | 180 | 300 | 420 |
|---|---|---|---|---|---|---|---|---|---|---|---|---|---|---|
| Grb2 + LATV3, full stimulus | Grb2, full stimulus | any | | | 0.01 | 0.000 | 0.003 | 0.001 | 0.000 | 0.02 | | | | |
| | | central | | | | | | | | (0.07) | (0.06) | 0.003 | | |
| Grb2 + LATV3, Itk ko, full stim. | Grb2, Itk ko, full stim | any | | 0.03 | | | | | 0.02 | 0.03 | | | | |
| | | central | (0.06) | | | | 0.04 | (0.07) | 0.001 | 0.001 | 0.01 | | 0.009 | |
| Grb2 + LATV3, anti-B7 | Grb2, anti-B7 | any | | 0.002 | 0.000 | 0.02 | 0.02 | | 0.009 | | (0.06) | (0.06) | 0.04 | (0.06) |
| | | central | 0.05 | 0.05 | | | | | | | | 0.04 | | |
| Lck + LATV3, full stimulus | Lck, full stimulus | any | | 0.04 | | 0.02 | 0.04 | 0.001 | 0.001 | 0.001 | 0.000 | 0.000 | 0.000 | 0.001 |
| | | central | | 0.03 | | 0.000 | 0.001 | 0.000 | 0.008 | 0.008 | 0.05 | (0.06) | 0.007 | |
| Lck + LATV3, Itk ko, full stim. | Lck, Itk ko, full stim | any | | | | 0.02 | 0.000 | 0.001 | 0.04 | 0.02 | | | | 0.02 |
| | | central | | | | | 0.007 | 0.003 | (0.07) | | | | | |
| Lck + LATV3, anti-B7 | Lck, anti-B7 | any | 0.05 | | | | | | | | | | | |
| | | central | 0.05 | (0.06) | | 0.03 | | | | | | | | |
| Vav1 + LATV3, full stimulus | Vav1, full stimulus | any | | 0.000 | 0.000 | 0.000 | 0.006 | 0.009 | 0.03 | 0.001 | | 0.02 | | |
| | | central | | 0.002 | 0.002 | 0.02 | | | | | | | | |
| Vav1 + LATV3, Itk ko, full stim. | Vav1, Itk ko, full stim | any | (0.06) | | | | | | | | | | | |
| | | central | | | | (0.06) | | | | | | | | |
| Vav1 + LATV3, anti-B7 | Vav1, anti-B7 | any | | 0.000 | 0.02 | | 0.05 | 0.03 | 0.03 | | | | (0.06) | |
| | | central | | 0.01 | | 0.02 | | | | | | | | |

### Table S5 – to Fig. 6

#### To Fig. 6 panel B

| Condition | Comparison | Pattern | -40 | -20 | 0 | 20 | 40 | 60 | 80 | 100 | 120 | 180 | 300 | 420 |
|---|---|---|---|---|---|---|---|---|---|---|---|---|---|---|
| SLP-76, anti-B7 | SLP-76, full stimulus | any | | | | 0.005 | 0.000 | 0.000 | 0.000 | 0.000 | 0.001 | 0.002 | | |
| | | central | | | 0.01 | 0.005 | 0.02 | 0.005 | | | | | | 0.01 |
| SLP-76, Itk ko, full stim. | SLP-76, full stimulus | any | | 0.05 | | 0.001 | 0.000 | 0.003 | 0.000 | 0.000 | 0.000 | 0.006 | | |
| | | central | | | 0.000 | 0.002 | 0.002 | 0.003 | 0.04 | 0.04 | 0.05 | | | |
| SLP-76, Itk ko, anti-B7 | SLP-76, full stimulus | any | | | 0.03 | 0.003 | 0.001 | 0.001 | 0.008 | 0.003 | | | | |
| | | central | | | 0.000 | 0.006 | | | | 0.04 | | | | |

#### To Fig. 6 panel C

##### SLP-76 V3

| Condition | Comparison | Pattern | -40 | -20 | 0 | 20 | 40 | 60 | 80 | 100 | 120 | 180 | 300 | 420 |
|---|---|---|---|---|---|---|---|---|---|---|---|---|---|---|
| **Restoration to SLP-76 full stimulus** | | | | | | | | | | | | | | |
| SLP-76 V3, full stimulus | SLP-76, full stimulus | any | | 0.04 | | 0.03 | | | | | | | | |
| | | central | | | | | | | | | | | | |
| SLP-76 V3, Itk ko, full stimulus | SLP-76, full stimulus | any | 0.01 | 0.000 | 0.01 | | | | | | 0.05 | | 0.03 | 0.003 |
| | | central | | 0.03 | 0.03 | 0.04 | 0.02 | | | | | | | |
| SLP-76 V3, anti-B7 | SLP-76, full stimulus | any | | | | | | | 0.000 | 0.000 | 0.000 | 0.005 | | |
| | | central | | | | | | | | 0.05 | | | | |
| SLP-76 V3, Itk ko anti-B7 | SLP-76, full stimulus | any | | | | | 0.03 | 0.05 | 0.000 | 0.001 | | 0.02 | | |
| | | central | | | 0.02 | | 0.04 | | | | | | | |
| **Enhancement under matched stimuli** | | | | | | | | | | | | | | |
| SLP-76 V3, Itk ko, full stimulus | SLP-76, Itk ko, full stimulus | any | | 0.000 | 0.000 | 0.000 | 0.000 | 0.000 | 0.000 | 0.000 | 0.000 | 0.000 | 0.001 | 0.000 |
| | | central | | | | | 0.05 | | | | | | | |
| SLP-76 V3, anti-B7 | SLP-76, anti-B7 | any | | | 0.008 | 0.008 | 0.005 | | | | 0.02 | | | |
| | | central | | | 0.002 | | 0.03 | | | | | | | |
| SLP-76 V3, Itk ko anti-B7 | SLP-76, Itk ko, anti-B7 | any | 0.05 | | | | | | | | | | | |
| | | central | | | | 0.03 | 0.004 | | | | | | | |

##### SLP-76 Vav

| Condition | Comparison | Pattern | -40 | -20 | 0 | 20 | 40 | 60 | 80 | 100 | 120 | 180 | 300 | 420 |
|---|---|---|---|---|---|---|---|---|---|---|---|---|---|---|
| **Restoration to SLP-76 full stimulus** | | | | | | | | | | | | | | |
| SLP-76 Vav, full stimulus | SLP-76, full stimulus | any | | | | | | | | | | | | |
| | | central | | | | | | | | | | | | |
| SLP-76 Vav, Itk ko, full stimulus | SLP-76, full stimulus | any | | 0.03 | | | | 0.03 | 0.01 | 0.02 | | | | |
| | | central | | 0.02 | | | 0.04 | | 0.006 | 0.05 | | | | |
| SLP-76 Vav, anti-B7 | SLP-76, full stimulus | any | | | 0.01 | | 0.04 | 0.003 | 0.000 | 0.000 | 0.001 | 0.000 | | |
| | | central | | | 0.04 | 0.006 | | | | | | | | 0.03 |
| SLP-76 Vav, Itk ko anti-B7 | SLP-76, full stimulus | any | | 0.03 | | | | 0.03 | 0.01 | 0.02 | | | | |
| | | central | | 0.02 | | | 0.04 | | 0.006 | 0.05 | | | | |
| **Enhancement under matched stimuli** | | | | | | | | | | | | | | |
| SLP-76 Vav, Itk ko, full stimulus | SLP-76, Itk ko, full stimulus | any | | | | | 0.02 | | | | 0.04 | | 0.02 | |
| | | central | | | 0.003 | | 0.007 | 0.03 | 0.006 | 0.004 | | | | |
| SLP-76 Vav, anti-B7 | SLP-76, anti-B7 | any | | | 0.000 | 0.001 | | | | | | | | |
| | | central | | | 0.000 | 0.000 | 0.02 | 0.01 | | | | | | |
| SLP-76 Vav, Itk ko anti-B7 | SLP-76, Itk ko, anti-B7 | any | 0.05 | | 0.007 | 0.02 | 0.03 | 0.01 | | | | | | |
| | | central | | | 0.02 | 0.02 | | | | | 0.05 | | | |

### Table S6 – to Fig. 7

#### To Fig. 7 panel B

| Condition | Comparison | Pattern | -40 | -20 | 0 | 20 | 40 | 60 | 80 | 100 | 120 | 180 | 300 | 420 |
|---|---|---|---|---|---|---|---|---|---|---|---|---|---|---|
| Grb2, anti-B7 | Grb2, full stimulus | any | | | | 0.001 | | 0.02 | 0.02 | | | | | |
| | | central | | | | 0.04 | | 0.05 | | | | | | |
| Grb2, Itk ko, full stim. | Grb2, full stimulus | any | 0.03 | | 0.001 | 0.000 | 0.000 | 0.000 | 0.000 | 0.008 | | | | |
| | | central | | | | | | | | | | 0.01 | | |
| Grb2, Itk ko, anti-B7 | Grb2, full stimulus | any | | | 0.002 | 0.000 | 0.000 | 0.002 | 0.006 | 0.03 | | | | |
| | | central | | | 0.02 | 0.003 | 0.05 | 0.006 | | | | | | |

#### To Fig. 7 panel C

##### Grb2 V3

| Condition | Comparison | Pattern | -40 | -20 | 0 | 20 | 40 | 60 | 80 | 100 | 120 | 180 | 300 | 420 |
|---|---|---|---|---|---|---|---|---|---|---|---|---|---|---|
| **Restoration to Grb2 full stimulus** | | | | | | | | | | | | | | |
| Grb2 V3, full stimulus | Grb2, full stimulus | any | | | | | | | | | | | | 0.02 |
| | | central | | | | 0.000 | | | | | | | | 0.05 |
| Grb2 V3, Itk ko, full stimulus | Grb2, full stimulus | any | | 0.02 | | 0.000 | 0.004 | 0.004 | 0.002 | | | | | 0.02 |
| | | central | | | | | | 0.006 | | | | | | |
| Grb2 V3, anti-B7 | Grb2, full stimulus | any | | | | | | | | | | | | 0.04 |
| | | central | | | | 0.001 | | | | 0.04 | | | | |
| Grb2 V3, Itk ko anti-B7 | Grb2, full stimulus | any | | | 0.000 | 0.000 | 0.000 | 0.000 | 0.000 | 0.001 | | | | |
| | | central | | | 0.01 | 0.01 | | 0.04 | | | | | | |
| **Enhancement under matched stimuli** | | | | | | | | | | | | | | |
| Grb2 V3, Itk ko, full stimulus | Grb2, Itk ko, full stimulus | any | | 0.01 | | | | | | | | | 0.03 | |
| | | central | | | | | | 0.03 | | | | | 0.05 | |
| Grb2 V3, anti-B7 | Grb2, anti-B7 | any | | | | | | | | | | | | |
| | | central | 0.05 | 0.05 | | | | | | 0.04 | | | | |
| Grb2 V3, Itk ko anti-B7 | Grb2, Itk ko, anti-B7 | any | 0.04 | 0.02 | 0.04 | | | 0.02 | 0.02 | | | | | |
| | | central | | | | | | | 0.04 | | | | | |

##### Grb2 Vav

| Condition | Comparison | Pattern | -40 | -20 | 0 | 20 | 40 | 60 | 80 | 100 | 120 | 180 | 300 | 420 |
|---|---|---|---|---|---|---|---|---|---|---|---|---|---|---|
| **Restoration to Grb2 full stimulus** | | | | | | | | | | | | | | |
| Grb2 Vav, full stimulus | Grb2, full stimulus | any | 0.05 | 0.000 | 0.000 | | 0.05 | | | 0.05 | 0.002 | 0.000 | 0.02 | 0.04 |
| | | central | | | 0.01 | 0.03 | | 0.04 | | | | | 0.05 | |
| | | periphery | | 0.000 | 0.000 | 0.000 | 0.000 | 0.000 | 0.000 | 0.000 | 0.000 | 0.000 | 0.000 | 0.005 |
| Grb2 Vav, Itk ko, full stimulus | Grb2, full stimulus | any | 0.03 | 0.000 | 0.001 | | 0.02 | 0.04 | 0.01 | 0.000 | 0.000 | 0.000 | 0.002 | |
| | | central | | | 0.02 | 0.001 | 0.03 | 0.01 | 0.03 | 0.04 | | | 0.05 | |
| | | periphery | | 0.000 | 0.000 | 0.000 | 0.000 | 0.000 | 0.000 | 0.000 | 0.000 | 0.000 | 0.007 | 0.04 |
| Grb2 Vav, anti-B7 | Grb2, full stimulus | any | 0.04 | 0.000 | 0.000 | | | | | | | | | |
| | | central | | | 0.01 | | | 0.05 | | | | | | |
| | | periphery | | 0.001 | 0.000 | 0.000 | 0.000 | 0.000 | 0.000 | 0.02 | 0.01 | | | |
| Grb2 Vav, Itk ko anti-B7 | Grb2, full stimulus | any | | 0.000 | 0.01 | | 0.05 | | 0.03 | 0.01 | 0.001 | 0.007 | | 0.04 |
| | | central | | 0.03 | 0.000 | 0.01 | 0.003 | | 0.02 | 0.04 | | 0.03 | 0.05 | 0.03 |
| | | periphery | | 0.002 | 0.000 | 0.000 | 0.000 | 0.000 | 0.000 | 0.000 | 0.000 | 0.000 | 0.04 | 0.03 |
| **Enhancement under matched stimuli** | | | | | | | | | | | | | | |
| Grb2 Vav, Itk ko, full stimulus | Grb2, Itk ko, full stimulus | any | | 0.000 | 0.000 | 0.000 | 0.000 | 0.000 | 0.000 | 0.000 | 0.000 | 0.000 | 0.000 | |
| | | central | | | | 0.02 | 0.02 | 0.05 | | 0.05 | 0.04 | 0.03 | 0.05 | 0.05 |
| | | periphery | | 0.000 | 0.000 | 0.000 | 0.000 | 0.000 | 0.000 | 0.000 | 0.000 | 0.000 | 0.003 | |
| Grb2 Vav, anti-B7 | Grb2, anti-B7 | any | | 0.002 | 0.000 | 0.000 | | | | | | | | |
| | | central | 0.05 | | | | | | | | | | | 0.05 |
| | | periphery | | 0.007 | 0.000 | 0.000 | 0.004 | 0.000 | 0.001 | 0.004 | 0.008 | | | |
| Grb2 Vav, Itk ko anti-B7 | Grb2, Itk ko, anti-B7 | any | | 0.03 | 0.000 | 0.000 | 0.000 | 0.000 | 0.000 | 0.000 | 0.004 | 0.01 | | |
| | | central | | 0.02 | 0.01 | 0.02 | 0.02 | 0.02 | 0.01 | 0.03 | 0.02 | 0.01 | 0.02 | 0.01 |
| | | periphery | | | 0.001 | 0.000 | 0.000 | 0.000 | 0.000 | 0.000 | 0.000 | 0.009 | | 0.04 |
