## Supplementary figures and images for "Transient protein accumulation at the center of the T cell antigen presenting cell interface drives efficient IL-2 secretion"

### Supplementary figures

##### Figure S1 – to Fig. 1

A

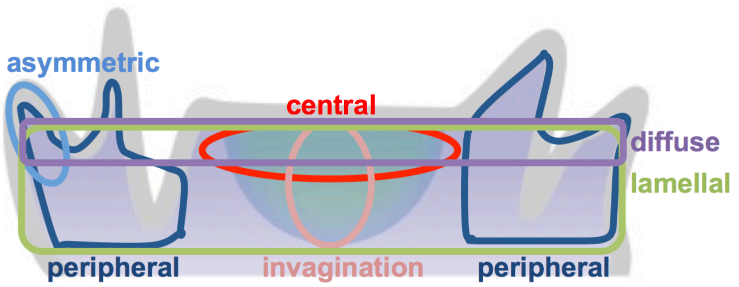


B

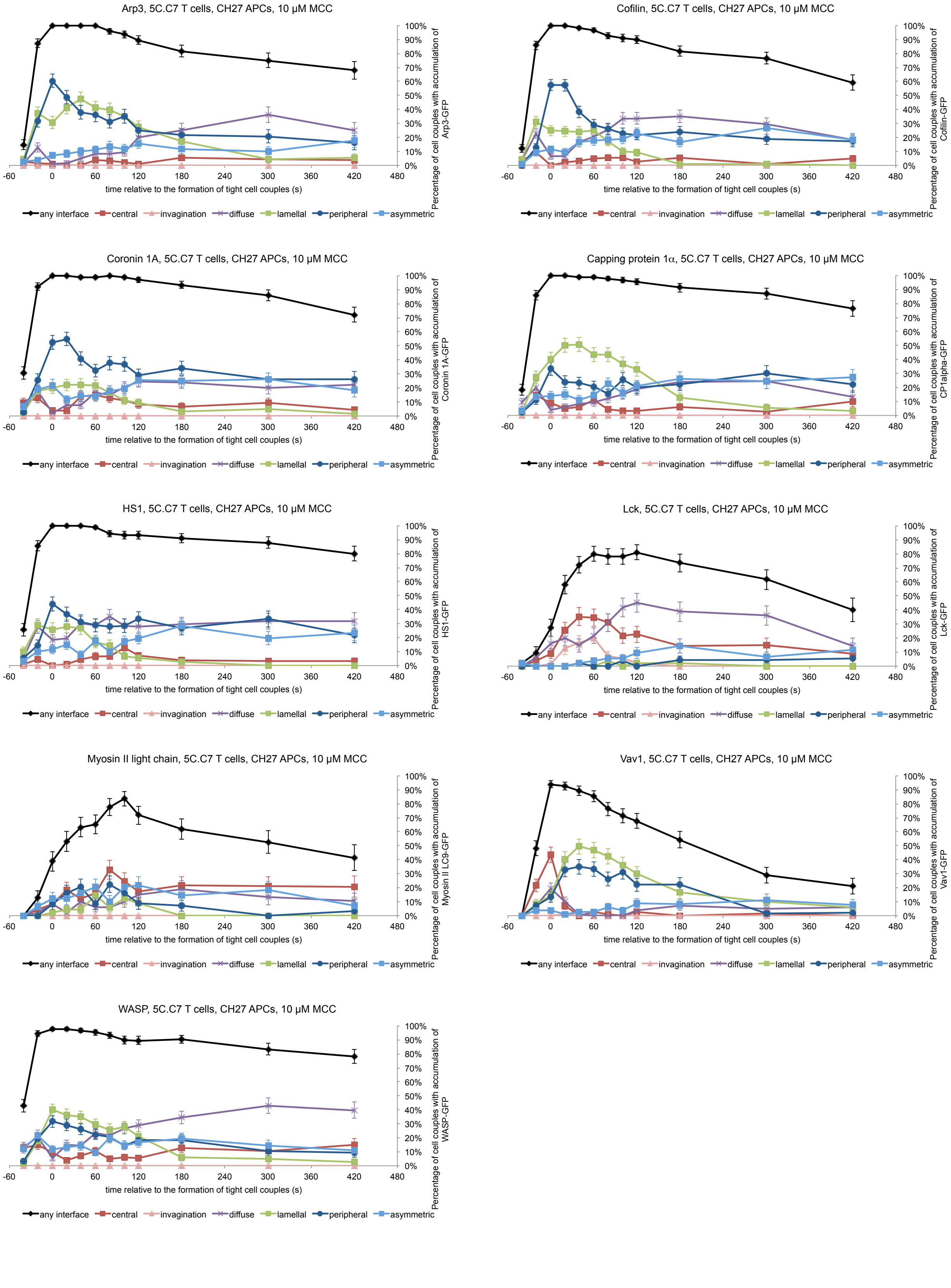

#### Figure S2 – to Fig. 1

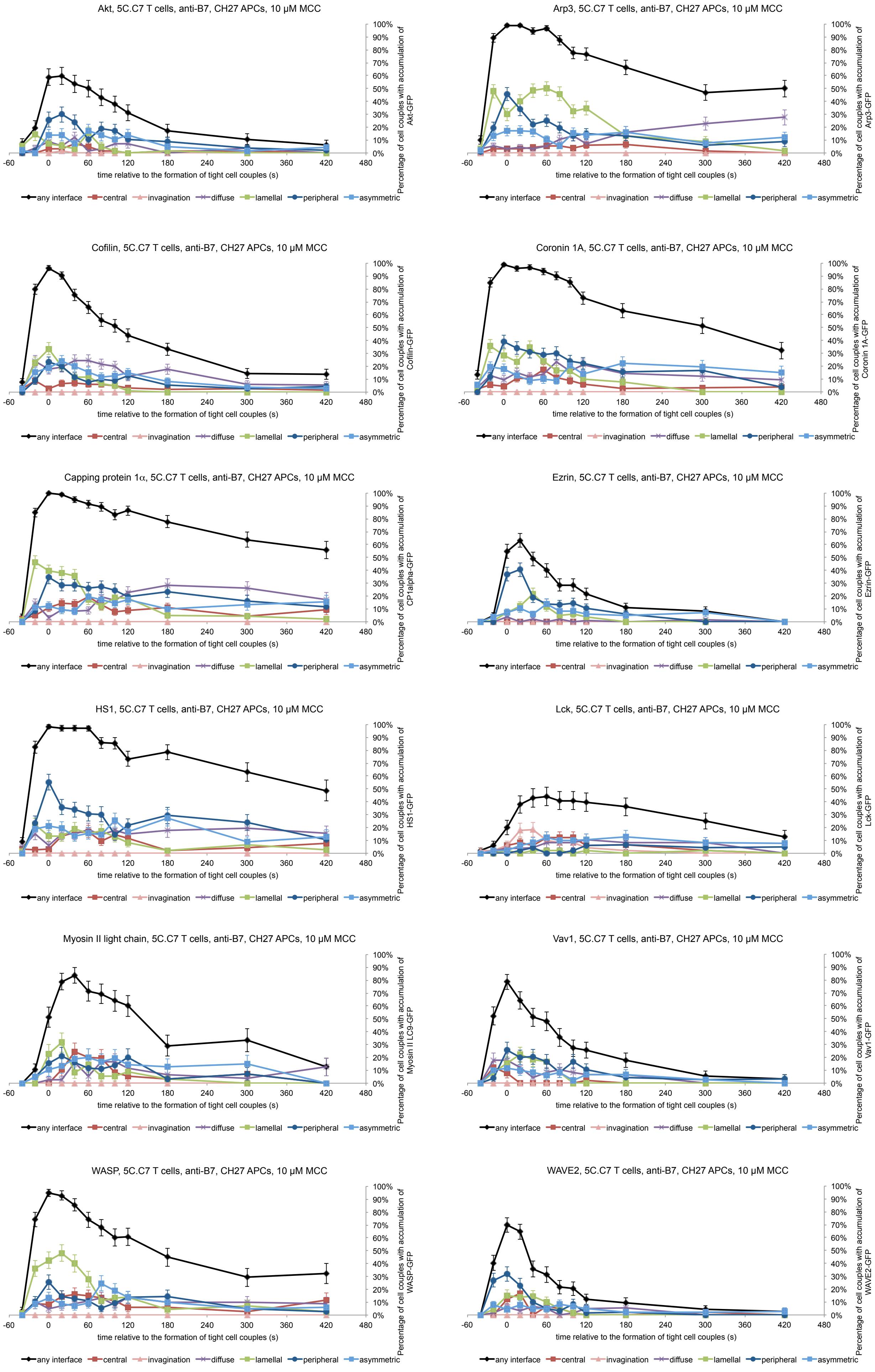

#### Figure S3 – to Fig. 1

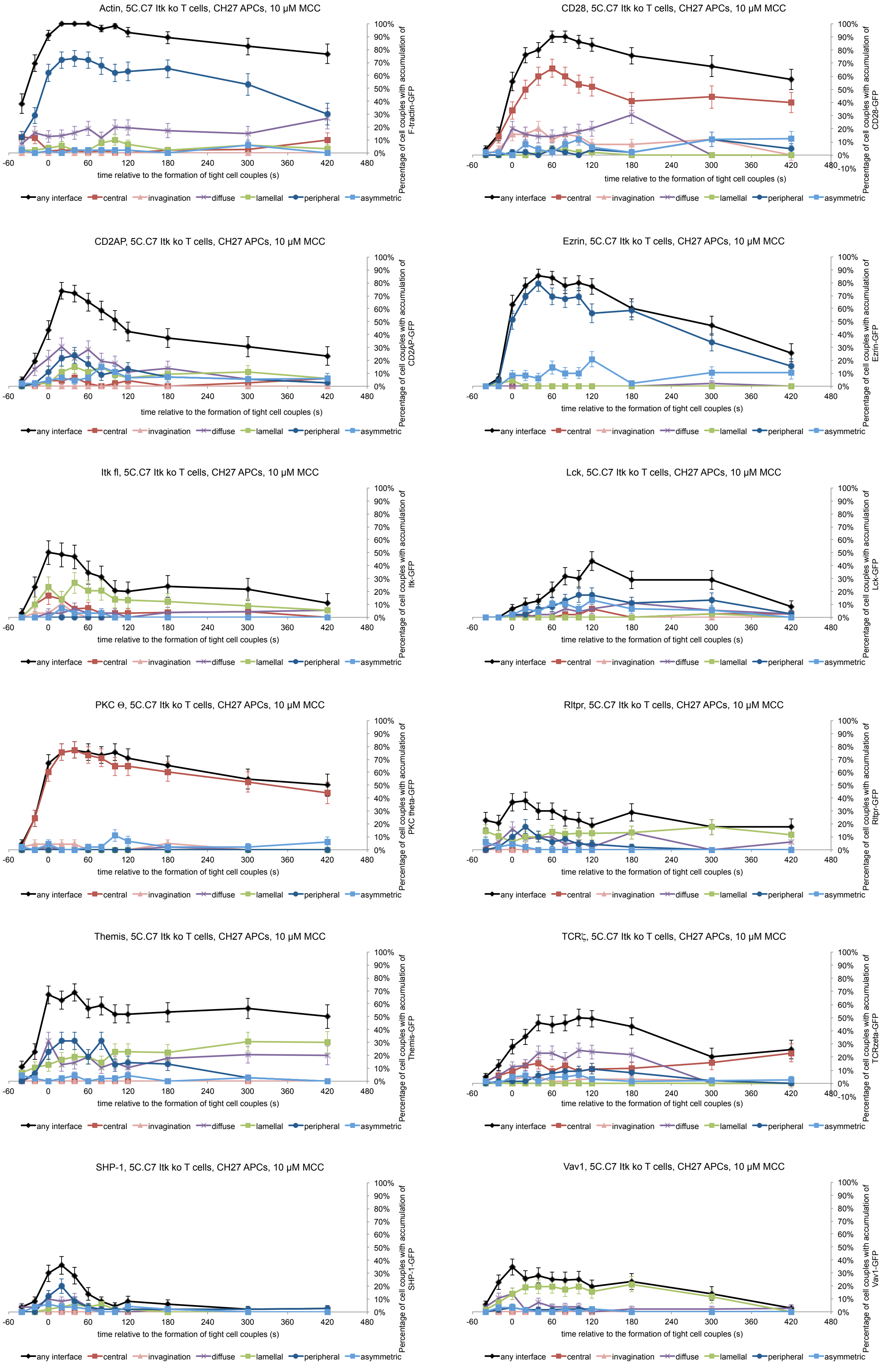

Figure S4 – to Fig. 2

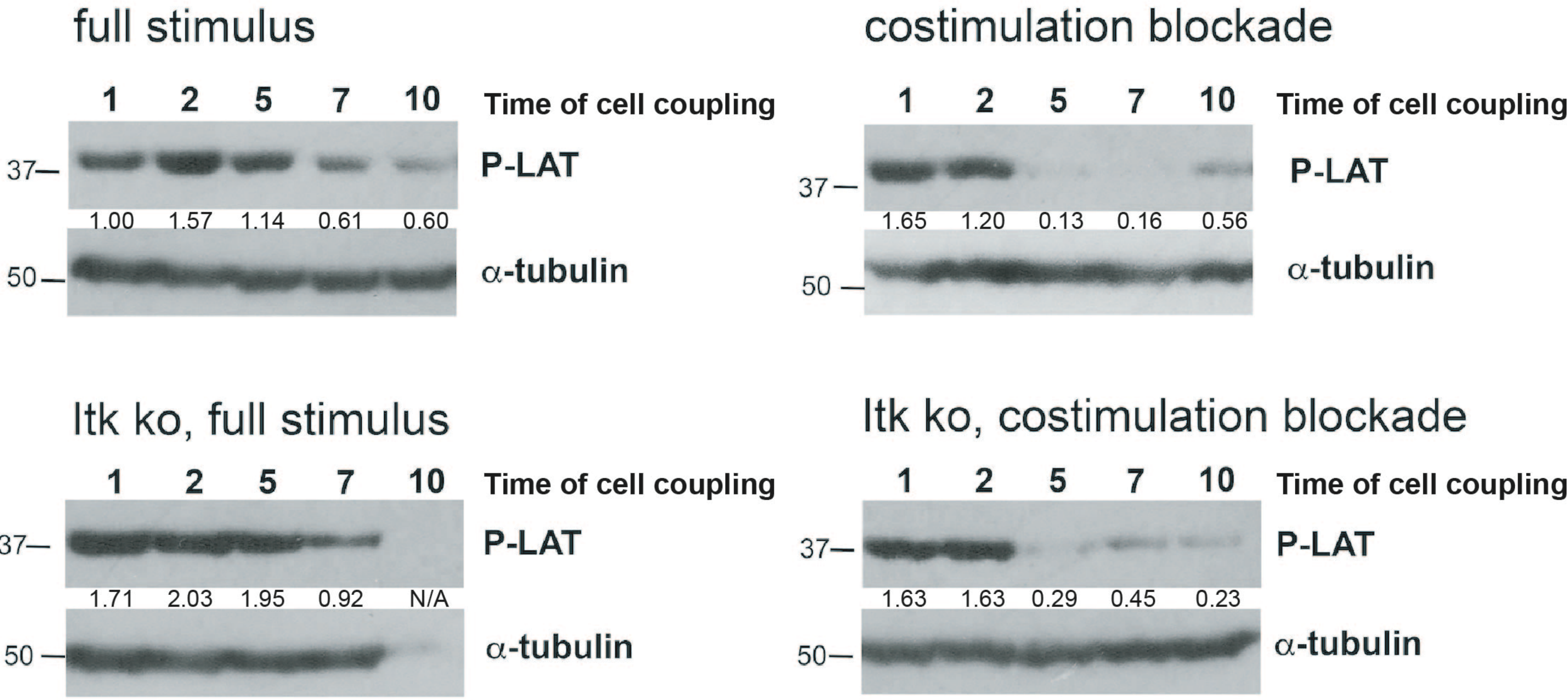

##### Figure S5 – to Fig. 4

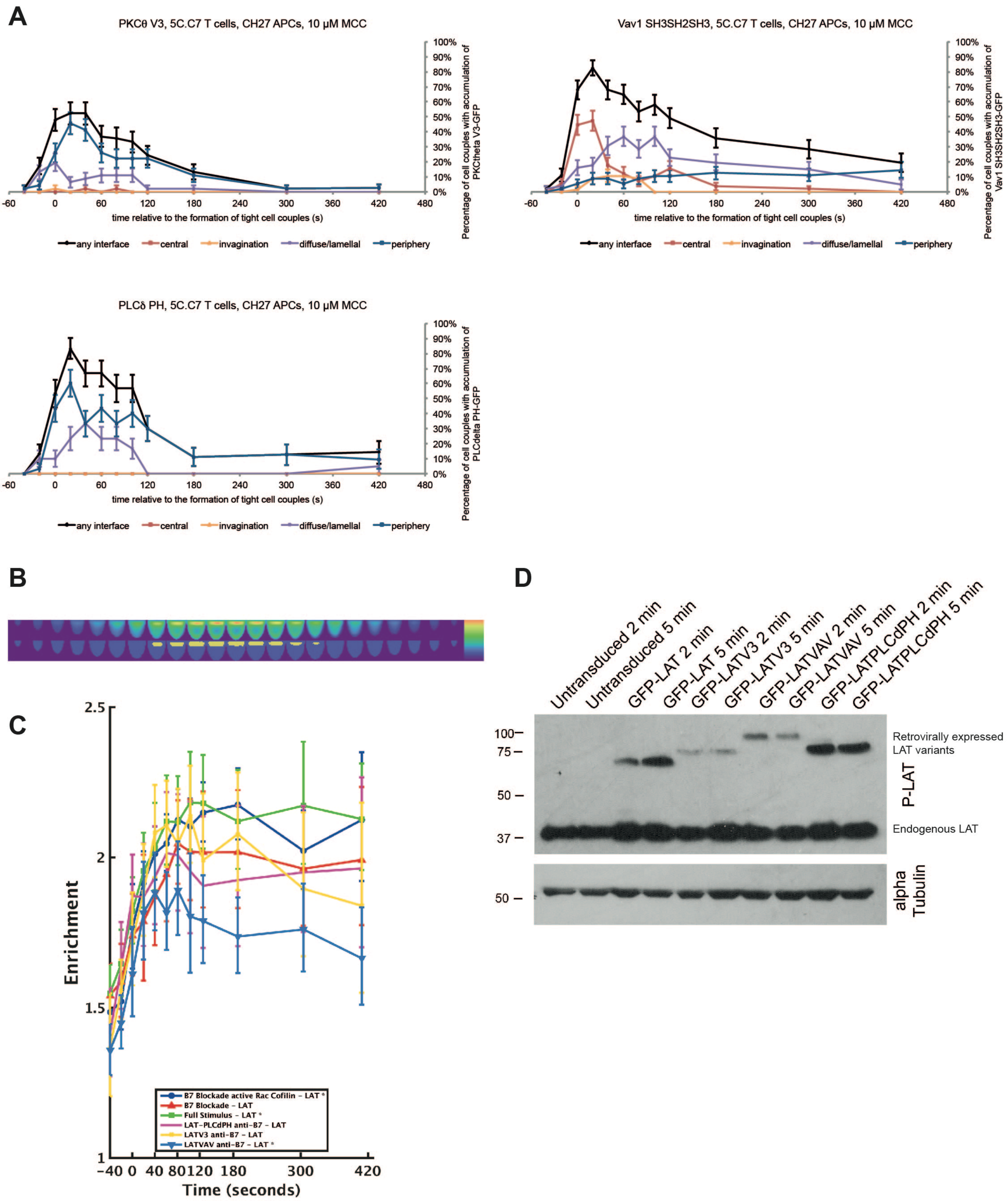

#### Figure S6 – to Fig. 5

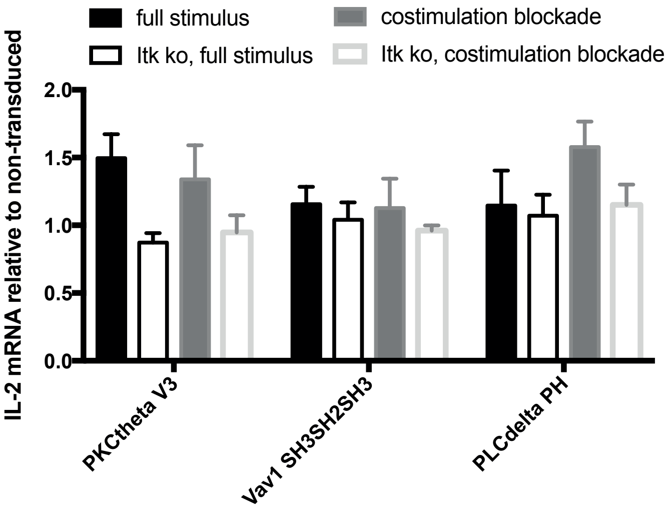
